# MultIscale MultiObjective Systems Analysis (MIMOSA): an advanced metabolic modeling framework for complex systems

**DOI:** 10.1101/718742

**Authors:** Joseph J. Gardner, Bri-Mathias S. Hodge, Nanette R. Boyle

## Abstract

In natural environments, cells live in complex communities and experience a high degree of heterogeneity internally and in the environment. Unfortunately, most of the metabolic modeling approaches that are currently used assume ideal conditions and that each cell is identical, limiting their application to pure cultures in well-mixed vessels. Here we describe our development of MultIscale MultiObjective Systems Analysis (MIMOSA), a metabolic modeling approach that can track individual cells in both space and time, track the diffusion of nutrients and light and the interaction of cells with each other and the environment. As a proof-of concept study, we used MIMOSA to model the growth of *Trichodesmium erythraeum*, a filamentous diazotrophic cyanobacterium which has cells with two distinct metabolic modes. The use of MIMOSA significantly improves our ability to predictively model metabolic changes and phenotype in more complex cell cultures.

## BACKGROUND

Microbes live in complex communities where they must interact with other organisms and compete for resources to thrive. By leveraging the capabilities of each individual in the community, consortia can achieve outcomes that are not possible by any one individual species. Metabolic engineers are learning from nature and are engineering synthetic consortia to take advantage of endogenous capabilities of specialists to achieve higher yields than pure cultures. One tool that has been used extensively to aide in the rational design of strains are metabolic models (11). The most widely used stoichiometric metabolic models are constraint-based linear programming models which vary in complexity from the relatively simple flux balance analysis (FBA) to more complex FBA models which integrate regulatory and/or thermodynamic constraints (12, 13) or time-dependent responses (14). The wide use of these models is due to the ease of constructing them; access to the genome sequence is enough to build a draft metabolic network. The simplicity of this technique does come at a cost: typical model formulations are limited to modeling steady-state growth of axenic cultures assuming homogenous environmental conditions; while this works for traditional metabolic engineering efforts on single species, it cannot accurately predict the behavior of consortia. There have been a few attempts to expand the applicability of these models to communities (3, 14, 15), but these models require assumptions that oversimplify the system, such as no diffusional limitations and identical or static growth rates for the different organisms. The current benchmarks for constraints-based metabolic modeling of microbial consortia are OptCom (16) and d-OptCom (17) (OptCom’s dynamic version). These approaches use inner and outer linear optimization problems to satisfy species and community level objectives, leveraging the inner solution as a constraint for the outer problem. However, this approach still relies on *a priori* determination of relative objective preference as well as predetermination of both species-level and community-level objectives. Additionally, cells are treated as homogenous spatial groups (13) or homogenous species groups (14), which limits the accurate simulation of cells acting individually, interacting with their environment, and ultimately forming communities. These approaches thus discount the complexity of individual cells forming communities and, instead of acting uniformly with neighbors or species, create dynamic intercellular and inter-environmental reactions (13, 18–20).

To more accurately model the complexity of community growth, a new modeling approach must be developed. We have developed <u>M</u>ult<u>I</u>scale <u>M</u>ulti<u>O</u>bjective <u>S</u>ystems <u>A</u>nalysis (MIMOSA), an advanced metabolic modeling framework for complex systems. This approach uses a multi-scale multi-paradigm metabolic modeling approach can leverage simple, powerful stoichiometric metabolic models and integrate spatio-temporal tracking of cells, nutrient diffusion, cell-cell interactions and cell-environmental interactions. This approach requires the use of both continuous and discrete variables as well as several different mathematical formalisms to reflect the multilevel behavior in populations. Therefore, we use an agent-based modeling (ABM) framework to allow direct interaction of different levels through the encapsulation of physiological, environmental, and metabolic models. ABM is a bottom-up modeling approach; the model is made up of a set of agents, which are allowed to act independently as long as they follow distinct rules of behavior defined by the user, this allows us to simulate emergent behavior of complex communities that arise from individual agent behaviors (21–24). The system behavior emerges as a result of the many (tens, hundreds, thousands, millions) individuals, each following their own behavior rules, living in a defined environment, interacting with each other and the environment (22). The integration of multiple modeling formalisms to represent disparate sub-systems is a trend common in engineering and science domains (25–29) and has recently seen some developments in the systems biology area (5, 30). Agent-based modeling has been previously applied to both intercellular (12, 31) and multi-cellular processes (32, 33) but has not previously been used to model metabolic fluxes. This multi-scale multi-paradigm approach represents a novel method of integrating individuals (through agents) with previously leveraged dFBA formulations (34, 35), thereby discretizing and separating variables for computational efficient solutions with low *a priori* knowledge.

As a proof-of-concept study, we chose to model *Trichodesmium erythraeum*, a filamentous diazotrophic cyanobacterium. *T. erythraeum* is a major contributor to the global nitrogen cycle; it is responsible for fixing an estimated 42% of all marine biological nitrogen (36) and it leaks 20-50% of the nitrogen it fixes (37), providing surrounding organisms with a biologically available nitrogen source. Unlike other diazotrophs, which either spatially or temporally separate the oxygen sensitive nitrogenase enzyme from the water splitting reaction of photosynthesis (oxygen production), *T. erythraeum* is unique because it simultaneously carries out nitrogen and carbon fixation during the day in different cells along the same filament (trichome). Therefore, it is the ideal model system for the development of MIMOSA: it has structurally identical cells that operate in two distinct metabolic modes (photoautotrophic and diazotrophic), a published genome scale model (3), transcriptome data, and a plethora of *in situ* and laboratory data to both train the model and validate predictions. We use this organism as a test-case for the modeling framework and illustrate how it can be used to develop a predictive model that can also be used to investigate cellular physiology by elucidating rules of behavior.

## RESULTS

### Model Formulation

We developed MIMOSA by integrating an updated version of the genome-scale metabolic model (3) (Table S1 for updated reactions) with nutrient diffusion, light diffusion, cell/cell interaction and cell/environment interactions (see Figure 1) using an agent based modeling framework. We have also implemented the use of multiobjective optimization to account for the dual cellular objective of producing biomass and the metabolite which is transacted between cells (glycogen or β-aspartyl arginine, depending on cell type). Constraints were imposed on the model as reported previously (3) with two notable exceptions. First, the ultimate product of nitrogen fixation was changed from ammonium to β-aspartyl arginine, which is the monomer used to create cyanophycin, a nitrogen storage polymer in *T. erythraeum* and other diazotrophic cyanobacteria (38–40). Second, the two major storage polymers, glycogen (modeled as maltose, or two linked glucoses) and cyanophycin (modeled as β-aspartyl arginine), were decoupled from the biomass formation equation so that they could freely accumulate or be metabolized. More detail about the formulation of the model is provided in Methods and Supplemental Text.

**Figure 1.**
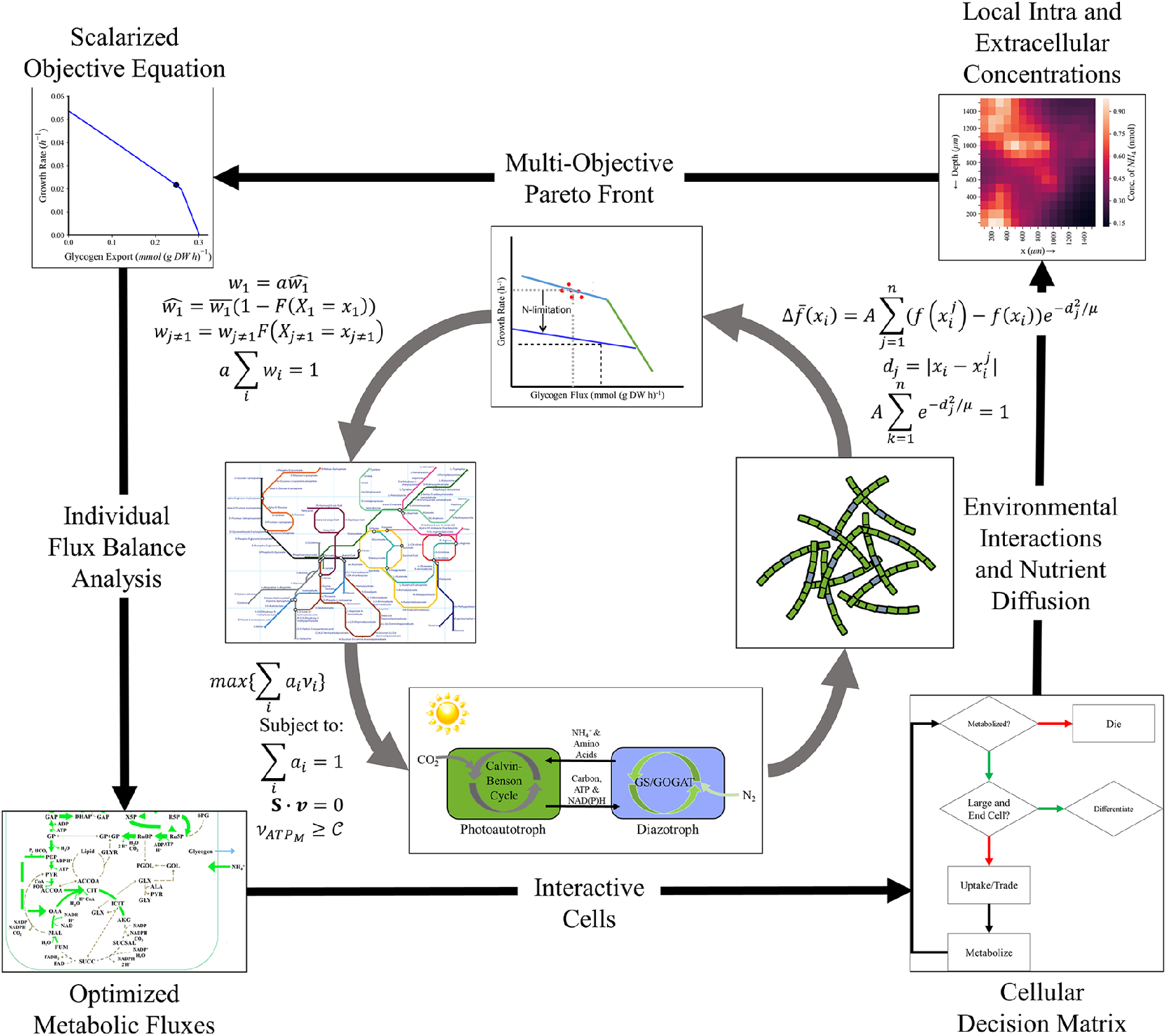
Multi-Scale Multi-Paradigm Model Generation. Before this process, the model generates an average scalar equation by fitting the organism’s Pareto Front to experimental data using the ATP hydrolysis maintenance reaction as further elucidated in Methods. Then, starting from the top and progressing with the arrows (clockwise): The multi-objective Pareto Front is corrected for environmental variables and cellular preferences using a weighting algorithm and assuming a normally distributed cell biomass (more detail in Methods). The corrected biomass equation is solved, individually, for each cell subject to existing constraints, a steady state over each time step, an appropriate maintenance ATP flux, and a scalar objective function for which all coefficients add to one. This is interpreted using the agent-based model to make individual cell and physiological decisions including 1) whether the cell should die, 2) whether the cell should reproduce (and if it does, what type of cell does it differentiate into), and 3) how it should interact with the environment and other cells. These interactions inform the status of the other cells (using an intrafilamental diffusion mechanism) and the environment (modeled with the same diffusion mechanism for CO_2_, N_2_, organic, and fixed nitrogen products, and assuming excesses of other media components). The iteration restarts with the objective equation updating each living cell (whether newly reproduced or previously established) based on the cell’s current metabolic state.

### Tracking Changing Cellular Objectives

MIMOSA evaluates the cellular objective for each cell for each time step based on the changing environmental conditions. As an example of this, we have tracked how the Pareto front changes for both photoautotrophic and diazotrophic cells over time (Figure 2). With increasing time, diazotrophs shift their objective away from biomass toward the production of cyanophycin as carbon becomes more available (Figure 2A). In contrast, photoautotrophic cells see a maximum production of glycogen at 9 hours after the onset of light and then their productivity decreases (Figure 2B). It is notable that every cell in the population is performing these decisions in parallel and Figure 2 is for a single representative cell of each cell type. Cell optimization changed based on environmental conditions and agent rules and the Pareto Fronts representing this behavior in these contexts is visualized in Fig. S1.

**Figure 2.**
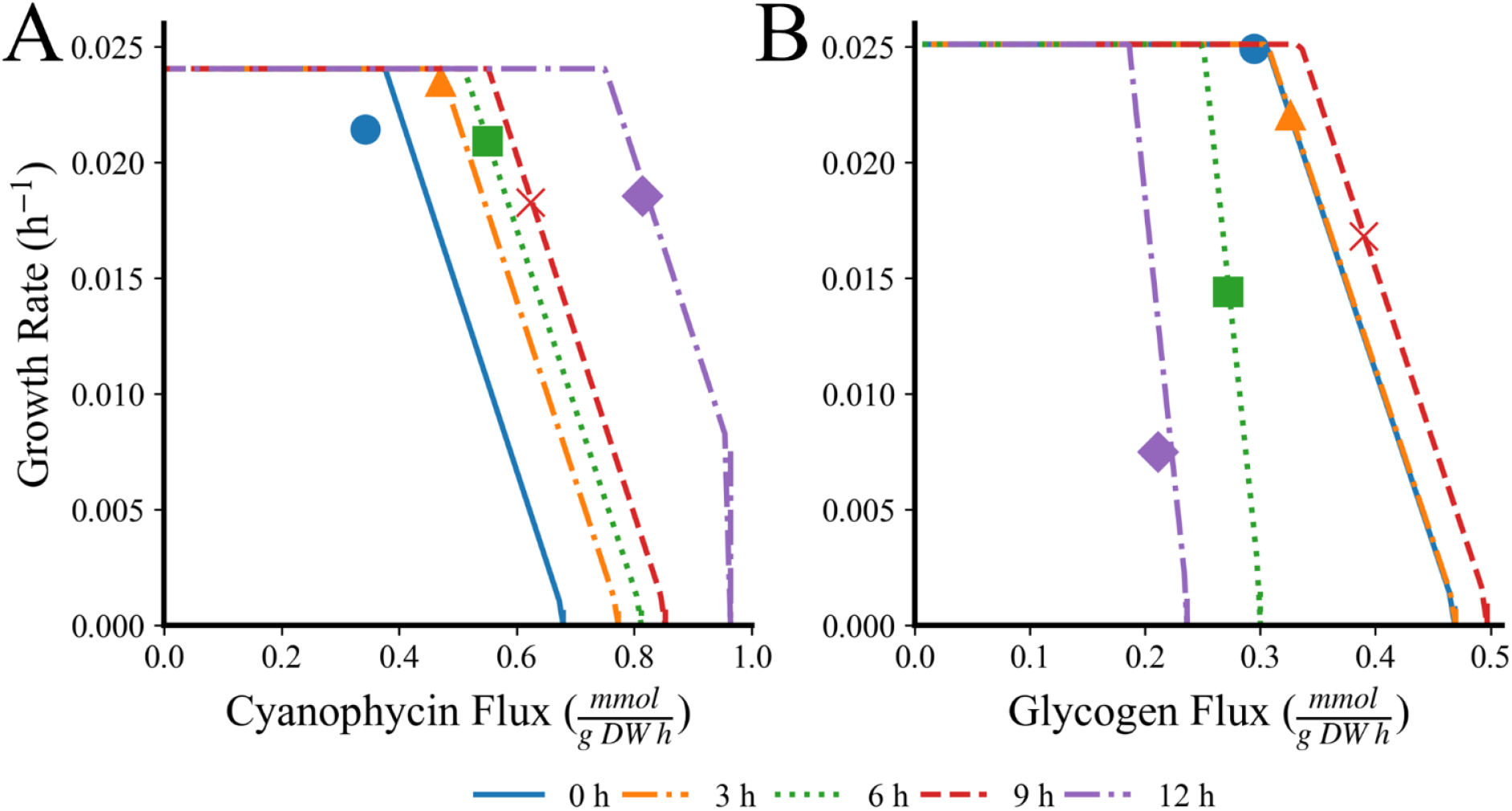
Pareto Front Progression and Selected Scalar Equation Points over the Light Period in Different Conditions for (A) a diazotrophic cell and (B) a photoautotrophic cell. Each line corresponds to a Pareto Front generated every 3 hours over the course of the light period at the start (blue, solid) through the end (purple, solid-dot) with the point selected by the simulation to best suit cell needs in the scalar objective function indicated. Simulations were run with atmospheric conditions, 100 *μ*E light, and YBC-II media with 150 cells over 10 filaments with a ratio of 4:11 diazotrophs to photoautotrophs.

### Model Validation

In order to test the predictive accuracy of the model, we predicted growth rate for a variety of different light intensities (Figure 3A) and compared to other published models for *T. erythraeum* (1, 2) as well as other experimentally measured growth rates (1, 3, 6, 9, 10, 41) exhibiting light saturation at higher light intensities. Ultimately, our model is a metabolic model, so it is important that it can also capture the metabolic changes that occur in response to changes in the environment. Therefore, we compared predictions of biomass changes to data collected in our laboratory for growth in different light intensities (see Figure 3B). The model was trained on data collected in 100 μE light and was validated with data collect in 50 μE light over a twelve-hour light period.

**Figure 3.**
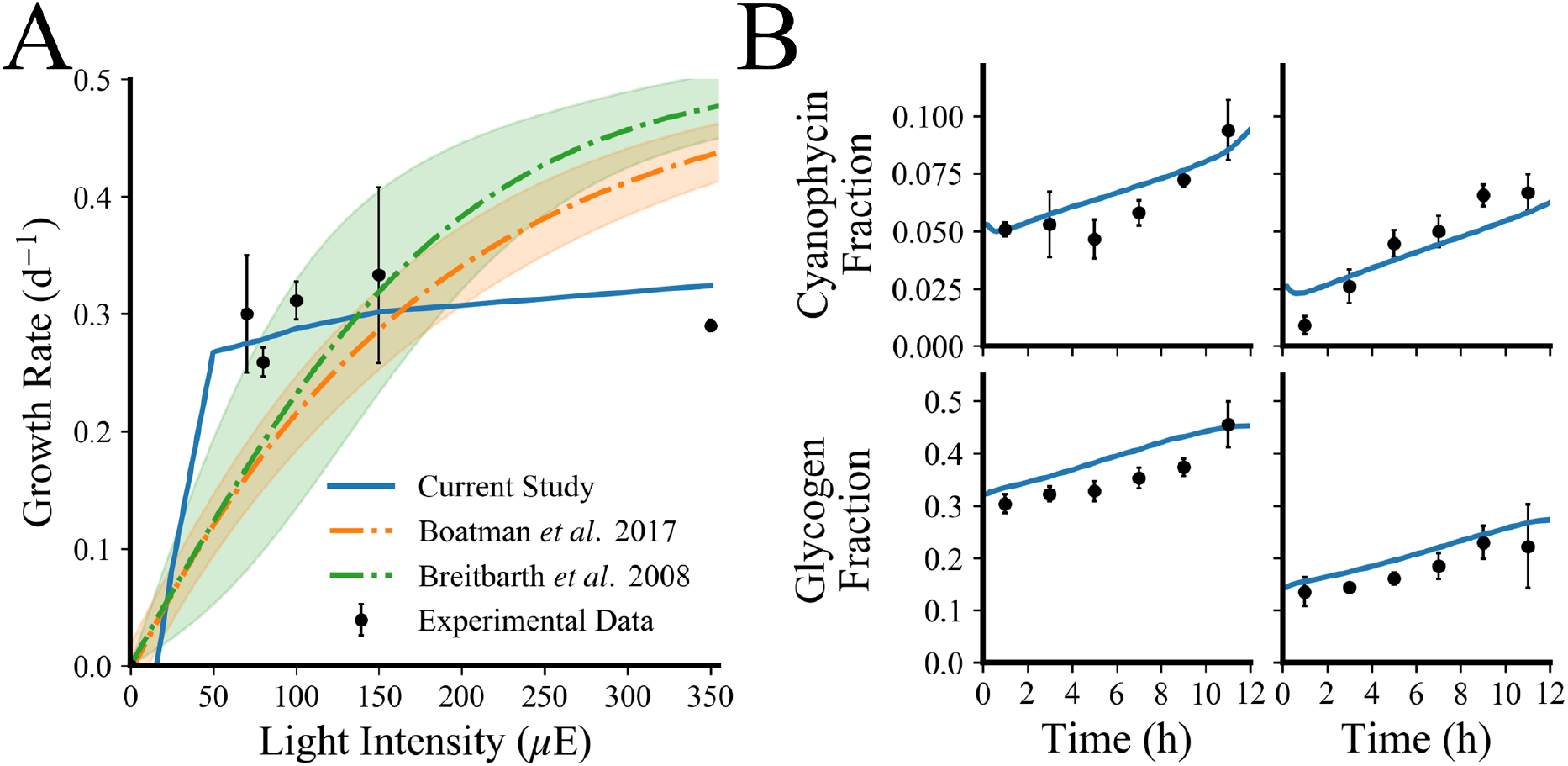
Simulation versus experimental data for model validation. (A) Growth rate as a function of light intensity in *T. erythraeum*; here we compare the predictions of growth rate from our model to two published models (1, 2) and experimental data reported from a variety of literature values (3–10). Error bars represent the error propagated when finding the mean of all separately recorded growth rates in similar conditions (YBC-II media, atmospheric CO_2_, no added nitrogen, and equivalent light intensity) using standard Euclidean error propagation. (B) Experimentally measured changes (black circles) in biomass accumulation for cells grown in 100 μE (B for cyanophycin and D for glycogen) and 50 μE (C for cyanophycin and E for glycogen) compared to simulated values (blue lines). Error bars on the glycogen and cyanophycin measurements represent one standard deviation for three biological replicates.

### Cells Alter Their Microenvironment

An advantage of the modeling approach we have developed is that we can track nutrients in the environment. Carbon dioxide (CO_2_) is typically the limiting substrate in aquatic photosynthetic growth due to low ambient concentrations and low solubility; for ambient CO_2_, Henry’s law defines an equilibrium concentration of 2.3 μM in the ocean. It is well known that photosynthetic microorganisms use carbon concentrating mechanisms (CCM) to concentrate CO_2_ near the carbon fixing enzyme, ribulose-1,5-bisphosphate carboxlyase/oxygenase (RuBisCO) to overcome low selectivity (42); our simulations imply that cells also increase the local concentration of CO_2_ immediately surrounding the cell (Figure 4A) and the release of nitrogen to the media including at more frequent time steps (Figure 4B). The simulation covers 150 cells and 10 filaments in a model 0.625 mm^3^ environment, corresponding to a filament density of 16 × 10^6^ trichomes m^-3^, well within the *in situ* ranges of free trichome density (43). This illustrates that the simulation corresponds well quantitatively to realistic local environments. At the end of our simulation, the cells on average can create a microenvironment that is roughly 2 fold higher in CO_2_ than the surrounding ocean. By looking at flux through major pathways, it appears that the CO_2_ is derived from high fluxes through the oxidative PPP and TCA Cycle in diazotrophic cells (Figure 5).

**Figure 4.**
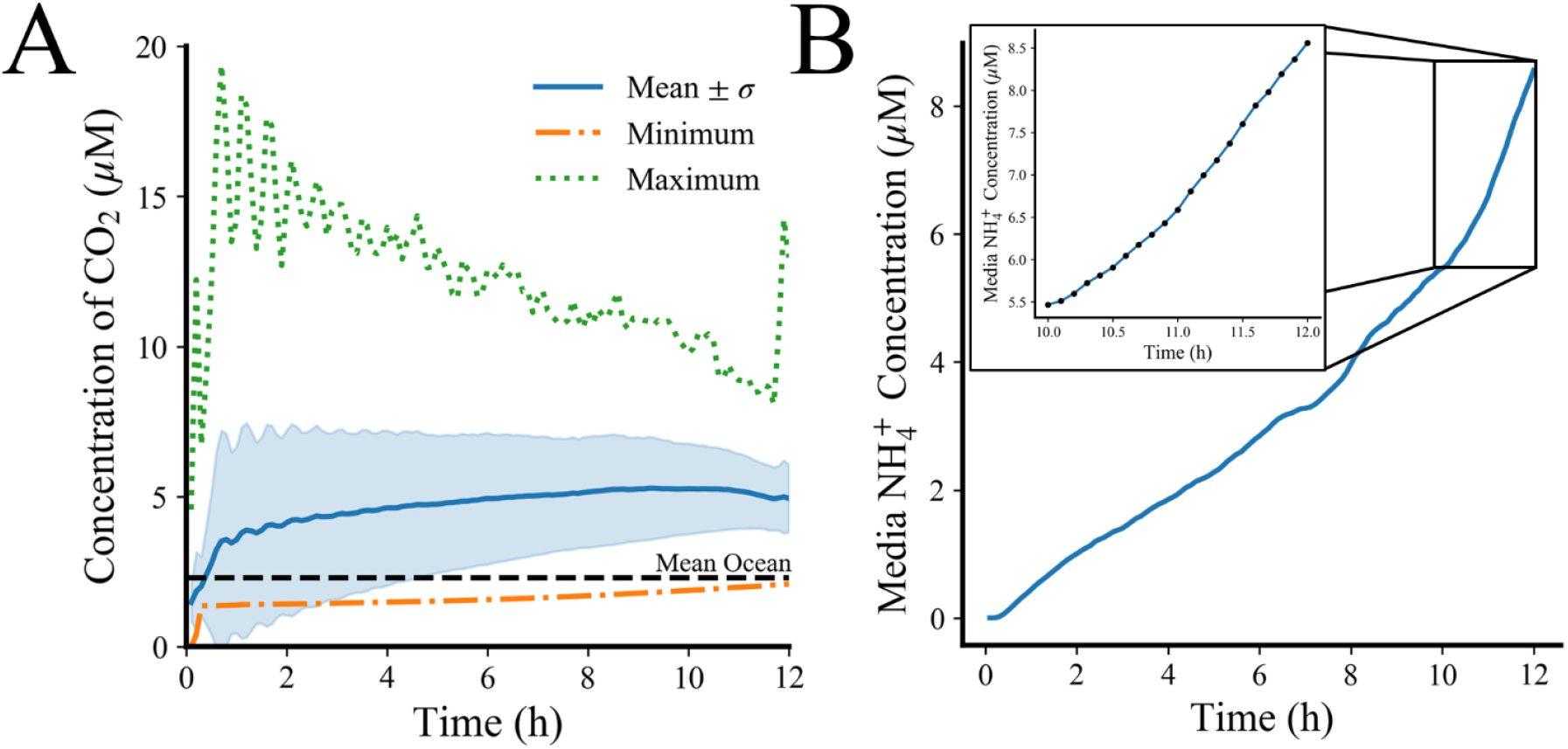
Cellular Interactions with the Local Environment. A) Local concentrations of CO_2_ in media surrounding cells. Blue line is mean ± 1 standard deviation, green line is maximum concentration in any ocean gridspace, orange is minimum concentration in any ocean gridspace, and the black line is the recorded mean oceanic concentration. 150 cells are present in the simulation in 625 square, 100*μ*m ocean gridcells with a maximum count of 25 cells gridcell^-1^ and a mean count of 0.302 ± 1.5 cells gridcell^-1^. B) Nitrogen release in a rough time step (0.5 hour) and a finer time step (0.1 hour) context allowing for investigation into more specific time periods while reducing computational load through state recapture.

**Figure 5.**
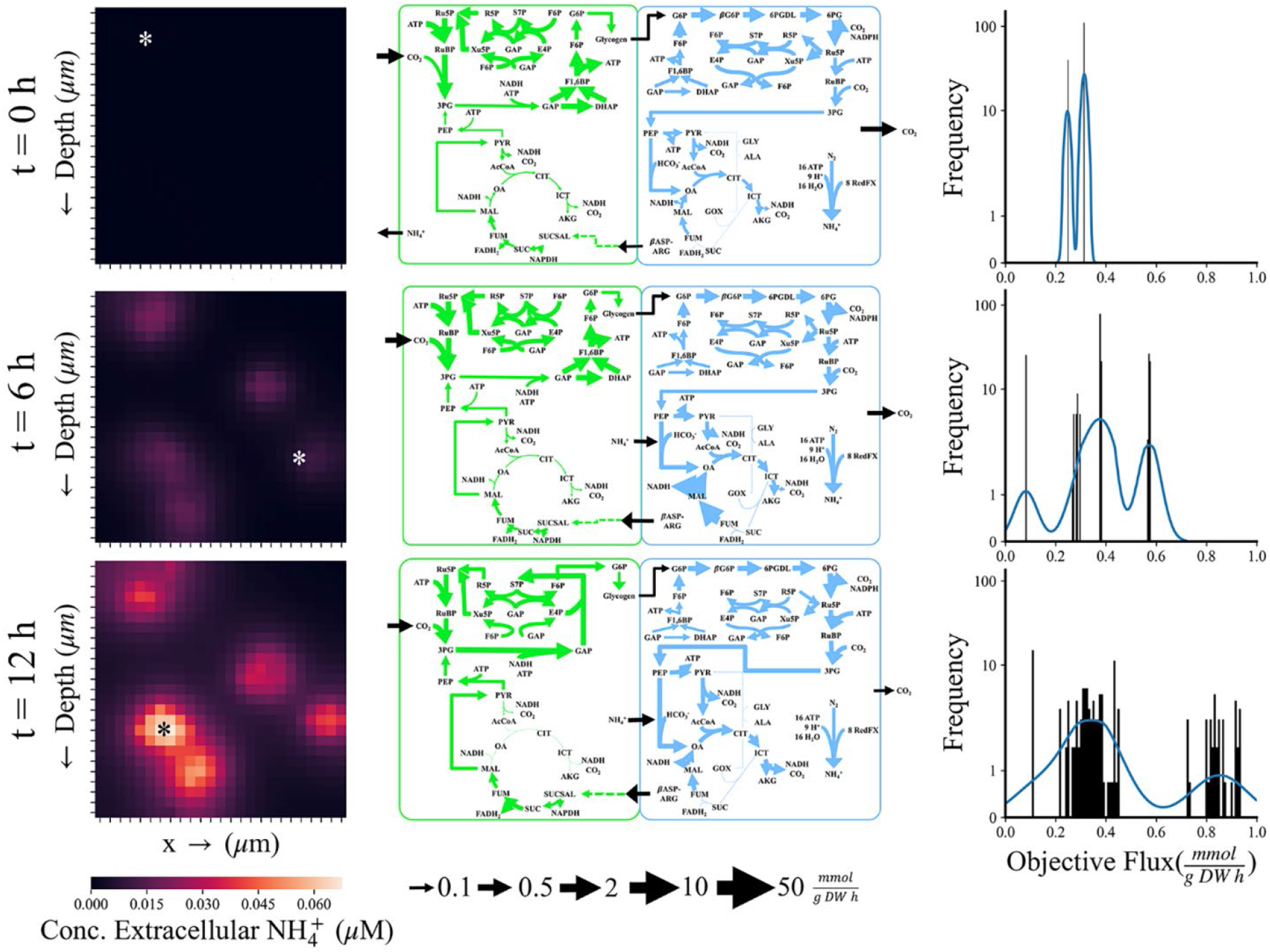
Examples of the type of data the multi-scale multi-paradigm metabolic model can track in time and space. Heat map legend and flux map legends are at the bottom of the figure. *x* ➔ indicates horizontal space in 2-D simulations. Time progresses from top to bottom with the top row indicating the beginning of the 12 hour light cycle cycle, the middle row the midpoint at 6 hours, and the bottom row the end of the light cycle. The left column contains the distribution of ammonium in the local media over these time points. The asterisk indicates the cells selected for the flux maps. The simulation can also be depicted as a 2-D grid with an arbitrary third dimension of one grid-cell deep. The whole space is a square grid with dimensions of 2500 *μ*m (2.5 mm) and each grid-cell with dimensions of 100 *μ*m (10 *T. erythraeum cells*). The middle column visualizes flux maps of photoautotrophs (left, green) and diazotrophs (right, blue) from the cells selected from the locations indicated by the asterisk. Flux maps were identical at 0 hours, selected from cells with the lowest extracellular ammonium in their gridspace at 6 hours, and selected from the highest extracellular ammonium in their gridspace at 12 hours. The right column is the frequency of unique metabolic profiles ranked by objective flux (a measure of cell productivity accounting for both biomass and metabolite production). Blue line is the kernel density estimate (kde) which estimates the probability of a given metabolic profile. Compound abbreviations for flux maps are provided in the “Abbreviations” section.

### Modeling a Heterogeneous Cell Population

One of the main advantages of this new modeling approach is that individual cells can be tracked in space and time so the heterogeneity of the population can be quantified (in terms of metabolic flux distributions). As an example, we tracked 150 cells over a 12-hour time period with time steps of 6 seconds which results in a total of 18,000 metabolic flux maps. Since this is an overwhelming amount of data to visualize, we have chosen to focus on a few representative flux maps (see Figure 5). In the left column, we track how the ammonium composition of the environment surrounding the cells changes with time from the initial seeding of cells at 0 hours to the middle of the daytime period (6 hours) to right before the onset of night (12 hours). These panels depict the release of ammonium into the environment as time progresses, and it is higher in areas where the cell density is highest. This agrees well with *in situ* data which reports that *T. erythraeum* leaks 30-50% of the nitrogen it fixes (37); our simulations predict that approximately 20% of the nitrogen fixed by the community is excreted into the medium. It is also important to note that the majority of ammonium is released by the cells in the second half of the day; during the first 6 hours, the cells release a total of 1.28 μmoles compared to 4.61 μmoles in the last six hours of the day. Again, this agrees with previous literature reports that the rate of nitrogen fixation peaks at midday (44), therefore we would expect more secretion of ammonium after peak nitrogenase activity. Select flux maps of cells growing in areas of low ammonium (top), medium ammonium (middle) and high ammonium (bottom) are depicted in the middle column of Figure 5. At the beginning of the simulations, cells are seeded in an environment that is identical to the defined marine medium YBC-II and because of this, they have identical flux maps as shown by the distribution graph in the right column. At time 0, we have a bimodal distribution because there are two cell types: photoautotrophic and diazotrophic. Photoautotrophic cells have high flux through the Calvin Cycle and the diazotrophic cells are operating in a more respiratory mode, with high flux through both the oxidative PPP and TCA Cycle. As the cells grow and start to experience more heterogeneity in their environment, they respond by differentiating their metabolism within the filament (Figure S2). First, this is evident in the frequency distribution plot, where they are both diverging in terms of total metabolic flux distributions and moving toward achieving optimal flux in terms of the objective function for both t = 6 hours and t = 12 hours. By comparing the changes that occurs in metabolic flux between areas of low, medium, and high ammonium, we can learn a few things about cellular physiology. In all cases, photoautotrophic cells have high flux through the Calvin cycle and an incomplete TCA Cycle, which has been widely reported in cyanobacteria grown phototrophically (45). In the case of *T. erythraeum*, succinic semialdehyde is derived from the nitrogen storage compound cyanophycin and is fed into the TCA Cycle to support the production of biomass precursors and glycogen (through gluconeogenesis). When external ammonium is high, photoautotrophic cells have less flux to glycogen, presumably because they do not need to provide as much to the diazotrophic cells to obtain fixed nitrogen in return. Investigations into imbalances in both metabolites and relative cell quantity display mechanisms of ammonium loss to the environment. Figure S3A illustrates how a lack of glycogen flux results in a higher loss of ammonium (with the exceptions of recently divided cells which metabolize glycogen with high ammonium loss) while Figure S3B visualizes a clear minimum ammonium release in the recorded range of percent diazotrophs per filament (between 15 and 30%). Diazotrophic cells have high flux through both the oxidative PPP and the TCA cycle which still utilizing carbon fixation reactions such as RuBisCO and PEP carboxylase and carbon conserving reactions like the glyoxylate shunt. Flux through the glyoxylate shunt increases as the availability of ammonium increases outside the cell, which is likely in response to the lower glycogen transfer from the photoautotrophs.

### Elucidating Rules of Cell Physiology

A key feature of agent-based modeling is the ability to model emergent behaviors in populations. We do not know all the rules of behaviors that define *T. erythraeum a priori* but by comparing simulations to observed *in situ* data and iterative improvement of the model, some rules can be elucidated. One trait that is widely variable in nature is filament length. It has been widely accepted that the average filament length is 100 cells (46) but more recent studies have suggested that they are typically much shorter, with a geometric mean of 13.2 ± 2.3 cells per filament, but with a mean range of 1.2 to 685 cells per filament *in situ* (47). Conditions for *in situ* sampling are widely variable so we hypothesized that filament length plays a role in maintaining growth in different environments: low light, low CO_2_ and low N_2_. We used the model to investigate which conditions might favor shorter or longer filaments (Figure 6). For each simulation, 150 total cells were seeded but in different trichome lengths (10, 30, 75, and 150 cells/filament) with and a ratio of diazotrophs to photoautotrophs of 3:7. In terms of growth rate, across all conditions we tested the shorter filaments had faster growth. This implies that diffusional limitations of nutrients into the cell and metabolites within the filament between different cell types start to hamper growth rate at longer filament lengths. The relative decline in growth rate is less dramatic for 25 μE when comparing across filament length, but when compared to other light conditions, there is a dramatic drop in growth rate for shorter filaments at low light. This indicates that longer filaments are capable of compensating for less light better than shorter filaments, perhaps due to increased surface area. Next, we examined the effect of filament length on cyanophycin composition for the same growth conditions as above. In every condition except low nitrogen, filaments with 75 cells appear to have more cells with above average cyanophycin content than other filaments lengths. Smaller nitrogen compounds (NH_4_^+^, amino acids, urea, etc.) can theoretically be used to support growth, permitting cyanophycin to be a longer-term storage compound. This is a possible explanation for the increase of cyanophycin in longer filaments. As filaments are longer, diffusive limitations become more pronounced, meaning that nitrogen gradients will remain in nitrogen replete cells longer and will be remade into cyanophycin as opposed to being metabolized for growth. This makes intuitive sense: not only is there a final drop-off at 150 cells, the distribution of cyanophycin content within the cells becomes larger, suggesting that some cells are starved for nitrogen and some are nitrogen replete. It is probable that filaments have adapted to leverage diffusion to both sequester nitrogen and to mitigate futile cycling of carbon and nitrogen compounds when diatomic nitrogen is available. The pattern of cyanophycin content diverges for cells in nitrogen limited environments due to overall shortages of nitrogen within the filament.

**Figure 6.**
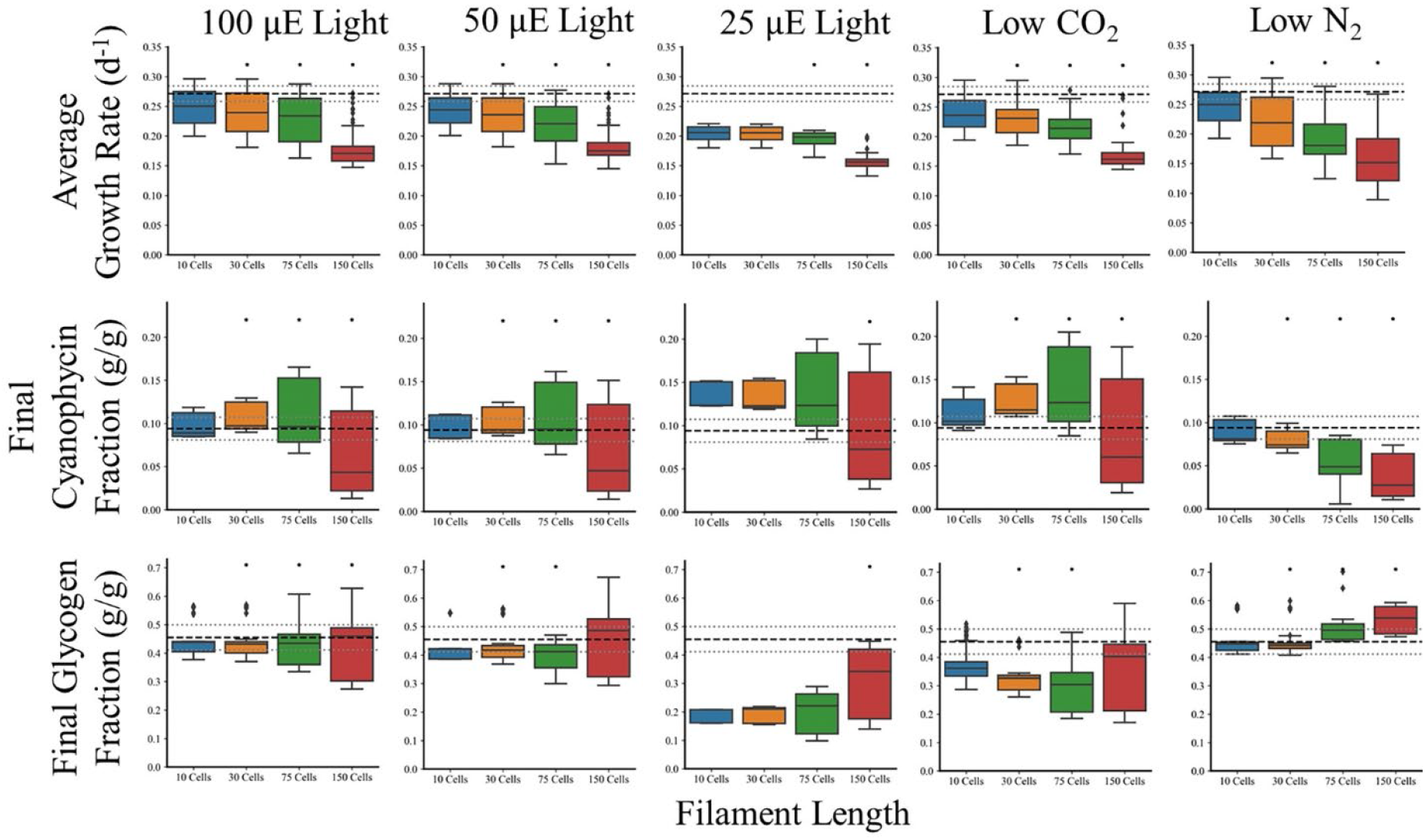
Trichome length affects the performance of the community. Trichomes were varied in initial length from short (10 cells filament^-1^) to long (150 cells filament^-1^) but with identical diazotroph: photoautotroph ratios of 3:7 (excepting 75 cells filament^-1^ which was 3.1:7) and initial cell counts (150 total) in the total population. Black asterisks are Student’s T-tests P-Values < 0.05 when comparing that cellular population to the 100 *μ*E case in every group. Dashed black lines are experimentally measured values and the gray dashed lines are their standard errors measured in the Boyle Laboratory (0.271 ± 0.0129 d^-1^ for growth rate, 0.0939 ± 0.0131 g cyanophycin/g DW, 0.455 ± 0.0440 g glycogen/g DW). Growth conditions represent the columns progressing as follows: 100 *μ*E light in YBC-II media and atmospheric conditions, 50 *μ*E light in YBC-II media and atmospheric conditions, 25 *μ*E light in YBC-II media and atmospheric conditions, 1.25 mM (versus 2.5 mM) HCO_3_^-^ and 200 ppm (versus 400 ppm) CO_2_ in otherwise atmospheric conditions and YBC-II media with 100 *μ*E light, and 0.395 atm (versus 0.79 atm) N_2_ in otherwise atmospheric conditions and YBC-II media with 100 *μ*E.

Finally, we investigated how glycogen content of cells changes due to filament length. The first pattern to note is that as length increases, the heterogeneity of the filament in terms of glycogen content also increases. This illustrates the importance of tracking individual cells because they are experiencing different environments and responding in different ways. Longer filaments also appear to be able to maintain glycogen content more readily than shorter filaments in all stress conditions we tested. Finally, nitrogen limited growth results in increased glycogen content as seen in other cyanobacteria (48). It appears that longer filaments in N limited growth can accumulate more carbon, perhaps again due to higher surface area and hence more energy from light harvesting. Our simulations agree well with published studies; it has been reported that growth rate and light intensity are both inversely correlated to filament length (49). This data indicates that filament length is largely determined by external cues rather than genetically.

## DISCUSSION

MIMOSA enables the most detailed and accurate metabolic modeling of complex systems to date by allowing coupling of several different mathematical formalisms describing natural phenomena, behavioral rules, and metabolism into a multi-scale multi-paradigm model. In constructing MIMOSA, we have added several features to enable us to more accurately predict phenotypes. A key feature of MIMOSA is the use of a multi-objective optimization approach. Unlike fast growing bacteria, which have successfully been modeled using a single objective function of maximum biomass (50), slow growing organisms have more complex objectives. In our simulations, *T. erythraeum* cells must achieve a delicate balance between biomass formation and the production of either glycogen or cyanophycin due to the symbiotic relationship between two cell types in the same filament. Photoautotrophs cannot function optimally without a biologically available form of nitrogen from the diazotrophs and the diazotrophs cannot support their metabolism without reduced carbon from the photoautotrophic cells. The use of multi-objective optimization allows us to describe this trade-off more accurately and by calculating the Pareto Front *a priori* we can also reduce computational effort. We have also accounted for changes in biomass composition that occur in response to changes in the environment or as a result of building carbon and nitrogen reserves during the day by decoupling the biomass equation. This allows the model to respond more fluidly to changes in the environment, which more closely mimics what cells experience in nature; for example, macro- and micro-nutrient stresses have been well known to cause changes in metabolism such as lipid and carbon accumulation (51–56). As such, the inclusion of metabolite and nutrient diffusion to augment metabolic optimization is a critical aspect of the model.

The influences of nutrient and energy availability in conjunction with population characteristics were studied to determine community and cellular adaptations to environmental perturbations. The model allows us to quantify the changes in the microenvironment around the cell compared to the bulk properties of the environment (Figure 4A) as well as to see how these changes affect the distribution of carbon and nitrogen inside the cell (Figure 5). These can be supplemented with “zooming in” on specific time steps to enhance investigation to rapidly occurring phenomena (Figure 4B). Not only did our predicted growth rates quantitatively match the experimental data, it was better able to capture effect of light saturation on growth rate; light intensities above 100 μE have little to no effect on growth rate (2, 49, 57–59). Our simulations agree well with the experimental data, however, there are differences that can be explained by the differences between our experimental conditions and our simulations. The main difference being the effect of diurnal light; *T. erythraeum* will not grow without diurnal day/night patterns, therefore the experimental data was collected from cells that were grown in 12 h: 12 h day/night cycles but the model is for a single 12-hour day time period. The addition of diurnal light patterns in future iterations of this model will help to improve the light dependent growth phenotype. Even so, the model is able to visualize community coordination and development during the 12 hour light period, exhibiting the increased release of ammonium to the media in the afternoon, consistent with the observation that nitrogenase activity peaks midday (44). Moreover, the individualized resolution of metabolic optimization can probe the nuances of intercellular, intracellular, and cell-environment interactions. Analysis of metabolic flux reveals a spontaneous partial/linear TCA Cycle in photoautotrophic cells consistent with previous reports (45). Cells also naturally coordinate to provide glycogen and cyanophycin transfer between cells, yielding oxidative behavior in diazotrophic cells through glycolysis with the possible side effect of oxygen consumption as a mechanism to protect nitrogenase as suggested in experimentation (58). Meanwhile, photoautotrophs naturally perform reductive carbon fixation coupled with utilization of the lower TCA Cycle to degrade arginine. These metabolic functions are affected by extracellular forces which are integrated into this model. For example, high ammonium environments result in declining gluconeogenesis in photoautotrophs (12 hours in Figure 5), likely since these cells are energetically limited and use cyanophycin as an energy source instead of light. Diazotrophs are prone to these environmental cues as well as low ammonium environments enhance light TCA Cycle to enhance recycling of amino acid byproducts from a lack of nitrogen. These observations, coupled with the diversity of metabolic profiles available to a relatively small population, By integrating modeling of other phenomenon with constraints based metabolic models, we were able to simulate *T. erythraeum* cultures that more accurately represent both *in situ* and laboratory data.

One of the many advantages of using this multi-paradigm framework is that we can simulate emergent behavior of a population. *In situ* data reports a wide mean range of trichome length from 1.2 to 685 cells (47); we used the model to investigate possible causes because this is a difficult phenotype to investigate experimentally. Our simulations suggest that even though longer filaments suffer from diffusional effects that limit growth, they are better able to handle stress (Figure 6) consistent with literature. Increased surface area in longer filaments minimizes the effect of lower light because the filament can harvest more light per volume. Also, the larger filaments are better able to maintain the average composition of storage compounds despite low carbon or low nitrogen conditions. Therefore, we would expect in areas of nutrient or light stress, the filament length would be longer.

One of the other unusual phenotypes of *Trichodesmium* that we were able to investigate using MIMOSA was leaking 30-50% of the nitrogen it fixes. Nitrogen fixation is an incredibly energy intensive process, costing the cell 8 ATP per ammonium, so it is not clear why *T. erythraeum* would excrete 30 – 50%. Despite using optimization to solve for fluxes, which should minimize energy losses, our simulations predict approximately 20% of the fixed nitrogen is excreted into the medium (Figure S4) which implies that this is a metabolically driven phenomenon. Further investigation has led us to develop three hypotheses on why this occurs: carbon limitation in diazotrophs, energy limitation in photoautotrophs, and imbalances between photoautotroph: diazotroph ratios. In the first case, photoautotrophs are unable to create glycogen chains and instead must start from a higher energy substrate than carbon dioxide (like succinic-semialdehyde) or must perform glycolysis on arginine derivatives to achieve energetic viability (Figure S3A). Second, population imbalances cause nitrogen to be produced faster than it can be anabolized into β-aspartyl-arginine chains and is released into the media, meaning there is an optimal ratio of cell types (Figure S3B). It is also possible that carbon limited diazotrophs are unable to manufacture full β-aspartyl-arginine chains and proton imbalances require ammonium release to the medium instead of passage to surrounding photoautotrophs.

MIMOSA enables the tracking of cellular-level environmental changes and the impact that they have on a metabolic model, opening the door to more accurate modeling of multi-cellular systems and the *in silico* investigation of the complex interactions between different cell types within an organism, and different species in a community. This is the first report of a metabolic model that integrates nutrient and light diffusion, cell/cell interactions and cell/environment interactions and we have used it to accurately predict growth, cellular composition and to investigate the unique physiology of *T. erythraeum*, which has filaments of both diazotrophs and photoautotrophs in close proximity. It establishes that this organism can effectively adapt to different conditions at three levels: the genetic level through division of labor in separate cell types, the metabolic level through relatively open-ended metabolic capabilities as well as further division within types, and at the population level to harness diffusional and physical interactions with the environment. MIMOSA is also a readily adaptable modeling framework – the addition of additional species to the model only requires the availability of a genome-scale metabolic and a few rules of behavior to be added. While we focused the proof-of-concept study of *T. erythraeum*, MIMOSA is a modeling framework that can be used to model a variety of more complex systems including applications in ecology, human health and metabolic engineering.

## MATERIALS/METHODS

### Cell Culture Conditions

Cells were grown as described previously (3). *Trichodesmium erythraeum* IMS101 cells were acquired from the Bigelow Laboratory for Ocean Sciences (East Boothbay, ME, USA). Cells were grown in a New Brunswick (Hamburg, Germany) with 100 and 50 μE in 12h light/12h dark cycles. Cells were grown in artificial seawater YBC-II medium (60) at pH 8.15-8.20. CO_2_ was maintained at atmospheric concentration. All chemicals were obtained from Sigma-Aldrich (St. Louis, MO). Growth rate was monitored by measuring chlorophyll absorbance (61) from 50 mL of culture every two days. Cyanophycin and glycogen were measured every four hours from the beginning of the light cycle (9 AM) to its end (9 PM). Total biomass mass was determined by dry weight analysis, cells were filtered with a Whatman 0.22 μm cellulose-nitrate filter and dried overnight at 100°C.

### Biomass Quantification

Carbohydrates were measured colorimetrically using the anthrone method (62) against glycogen as a standard. Cyanophycin was extracted by disrupting 740 μL of 250 mL cells concentrated to 2 mL via filtration and rinsing with TE buffer with 2.70 mg/mL lysozyme overnight at 37 °C, centrifuging at 16,100 x G for 5 min, and resuspending the pellet in 1 mL of 0.1 M HCl (in which cyanophycin is soluble) for 2 h. The extraction was repeated on the pellet, the supernatant fractions were combined, and cyanophycin was quantified colorimetrically using the Sakaguchi reaction (63).

### Mass Balance Constraints

Constraints based metabolic models are based on mass balances, therefore it is imperative that we develop accurate accounting of each element. Therefore, we used training data (Table S2) to estimate normal cellular consumption (Table S3). Average objective fluxes were estimated using mass balances around biomass and metabolite production with the formulation:

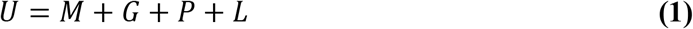

Where *U* corresponds to uptake, *M* is the nitrogen or carbon required for maintenance metabolism, *G* is the accumulation of fixed carbon or nitrogen during growth into non-biomass metabolites, *P* is the accumulation of fixed carbon or nitrogen that is passed to the other cells, and *L* is the carbon or nitrogen leaked into the surrounding media. This can be further detailed into carbon and nitrogen energy balances (defined as above with the subscript “N” for nitrogen and “C” for carbon):

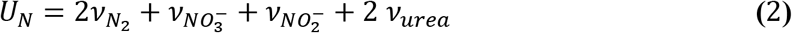

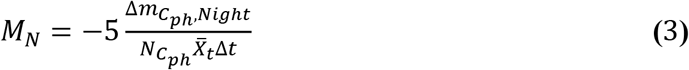

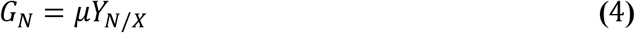

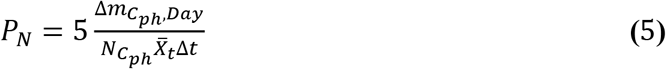

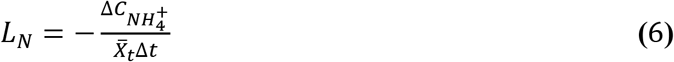

Where *ν* corresponds to flux of the substrate (indicated in the subscript), *m* is mass, *N* is molar mass, the subscript *c_ph_* is cyanophycin, 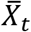 is average biomass over the measured time period (*Δt*), *μ* is growth rate, *Y_N/X_* is the nitrogen stoichiometry in biomass estimated by the biomass composition. The stoichiometric coefficients represent the number of nitrogen atoms in each molecule; 2 per diatomic nitrogen and 5 per *β*-aspartyl arginine. The flux of nitrogen (*ν*_*N*_2__) is measured and recorded via the acetylene assay for nitrogenase activity as recorded in the literature for the same growth conditions. *G_N_* was approximated using the model’s prediction for cellular composition of nitrogen using the biomass equation and balanced equations. The faction of cyanophycin in biomass was measured analytically at 6 timepoints throughout a single 12-hour daytime period (See *Biomass Quantitation* in methods). Ammonium release over a 12-hour period was below detectable limits (< 1 μg/L) (64) in our laboratory experiments. If we re-arrange this equation to solve for the average flux of nitrogen into a single cell, 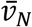, we obtain the following equation which can be used to solve for 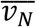 or *ν*_*N*_2__:

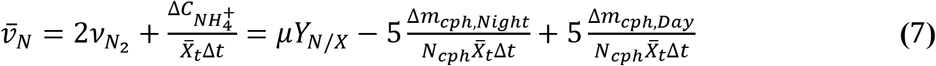

Assuming each cell requires the same amount of nitrogen, that only diazotrophs reduce diatomic nitrogen, that the average ratio is 4:1 photoautotrophs to diazotrophs (44) for estimation of training data for consumption and production, and that cells do not release ammonium at optimal production, maximum nitrogen flux into a photoautotrophic cell can be approximated as:

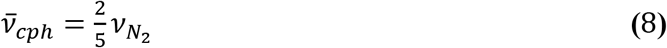

The same approach is taken for the carbon mass balance.

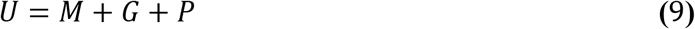

Where:

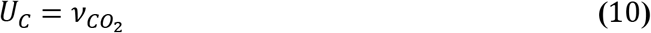

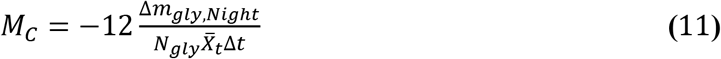

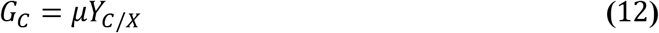

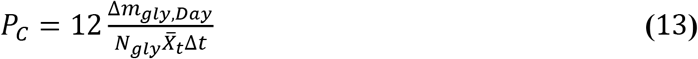

And:

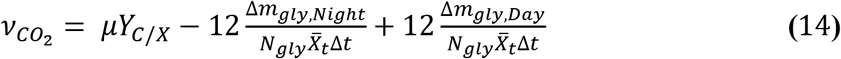

In this case, the variables are the same except for subscripts C (carbon), gly (glycogen). And CO_2_. *G_C_* represents the stoichiometric predictions of elemental composition and *ν*_*CO*_2__ is approximated using equation 10. This allows prediction of maximal glycogen flux (assuming 12 carbon molecules per glycogen, since it is modeled as disaccharide glucose or maltose) using:

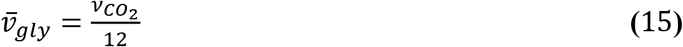

### Development of Agent Based Model

Repast Simphony (65) in Java was used as the agent-based modeling framework in which differentiated multi-objective metabolic models of *Trichodesmium erythraeum* are contained. It contains three agent types – Ocean, Cells, and Filaments. The Cells agent contains two sub-agents representing each cell type: photoautotrophs and diazotrophs and is responsible for intracellular processes and decisions. The Ocean agent defines and calculates the extracellular environment and the Filaments agent organizes the Cells and modulates their transactions.

#### cells

Cell agents (cells) are generated for each individual cell in the model. These contain two subtypes, photoautotrophs and diazotrophs, but contain several consistent elements between the two. Simulation variables are summarized in Table S4. All cells reproduce according to the same rules: cells divide according to sampling from the weighting distribution described above if that sample is bigger than the cell mean cell, cells only extend from the ends, and cells can only divide into diazotrophs if there is a diazocyte under development (decided at the filament level if the filament is nitrogen limited). When a cell is large enough, it converts to fully stationary growth, producing only metabolites and creating a larger and larger metabolic gradient between cells without *de novo* biomass synthesis. This prevents a cell from becoming excessively large in the center of the filament. Cells will die if they cannot produce the requisite maintenance ATP through metabolism or catabolism.

Cells allow metabolites to diffuse through the lipid bilayer using permeabilities reported in the literature (Table S5). This mechanism represents a non-zero leakage scenario that was nevertheless much slower than intrafilamental diffusion (Table S6). Scavenging from the environment for compounds which carried no evidence of active transport followed these same rules and was therefore prone to concentration gradients. Active transporters, on the other hand, allowed the cell to uptake whatever concentration of compound was necessary subject to its presence in the local ocean grid. Allowable exchange of metabolites between cells is illustrated in Figure 7. If several cells compete in that grid space, access to the available molecule was divided equally among those cells.

**Figure 7.**
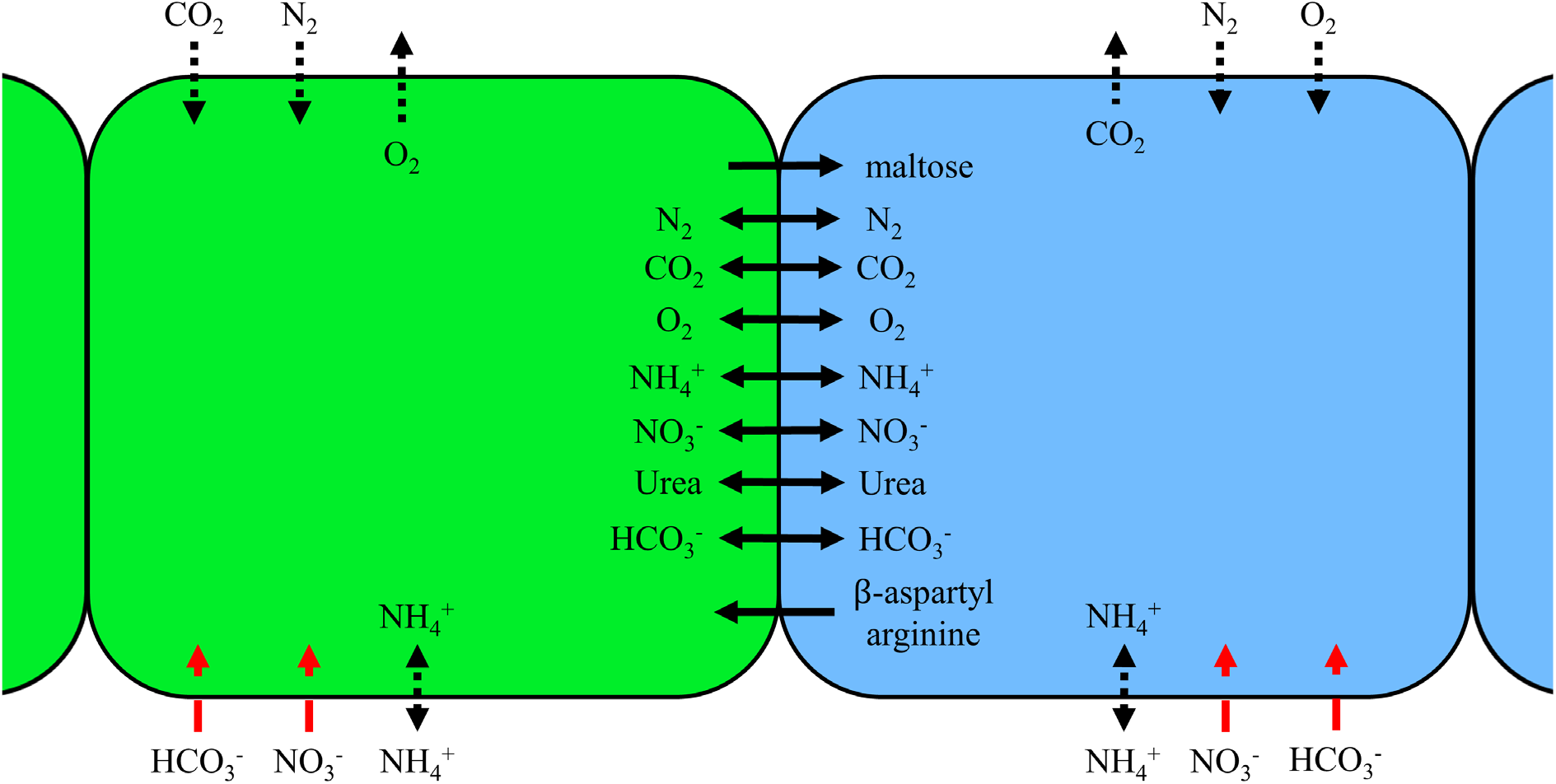
Allowed Metabolite Diffusion and Exchange between and into Cells. Green cells represent photoautotrophic cells and blue represent diazotrophic cells. Solid black lines indicate free diffusion between cells, dotted black lines indicate lipid bilayer diffusion, and red dashed lines indicate coupled ion transport into the cell. Not pictured is ATP synthase. Maltose is modeled as the 12-carbon molecule that forms the foundation for glycogen while *β*-aspartyl arginine is the foundation for cyanophycin.

#### Subclasses: Photoautotrophs and Diazotrophs

Both subclasses define the uptake constraints and send to a Python file that decides whether the cell metabolizes or catabolizes based on those constraints using the multi-objective metabolic model previously described (see *Supplemental Information: Routine Metabolic Optimizations*). The cell then updates its internal metabolites based on the optimization results, diffuses metabolites, divides if possible, and uptakes from its local environment. Model bounds are calculated using local concentrations to calculate maximum flux bounds excepting β-aspartyl arginine which is further limited to 8% of available nutrients (See Figure S5). These methods are handled by three ScheduledMethods that Repast Simphony schedules in specific progression. Together with the Ocean Agent’s updates, the individual cell actions (as dictated by the metabolic model) form the core of the simulation. A more detailed flow chart for cell decision making can be found in Figure S6. Progression through these steps is identical for cell types, but the metabolic static variables (objectives, gas uptake, etc.) are different between the two subclasses, necessitating separate methods.

#### Ocean

The ocean agent is responsible for tracking cells and modeling external nutrients. Its main task is facilitating diffusion between cells and locations as well as approximating an uptake radius for cells. Each ocean represents a uniform, static, abstract area of the overall grid space with a uniform dimension space of *δ* × *δ* where *δ* is a user defined parameter. This set of simulations was conducted with time steps of 0.1 hours as a moderate value between diffusion phenomena (on the order of seconds along the length of a filament) and doubling time (on the order of 50 hours). Metabolites are assumed to freely diffuse in a dilute seawater environment between cell filaments (Table S6) and assumed to be uniform within the grid, given the relatively long time step compared to the rate of diffusion over such small dimensions. If the impacts of metabolic diffusion limitations were of interest, the time step within the framework could be made appropriately small to more accurately track metabolites, at cost of increased computational burden. Each ocean gridcell diffuses molecules into its adjacent ocean gridcells assuming discretized slab diffusion in two dimensions. This is done using a previously developed discrete algorithm for diffusion in a grid (66):

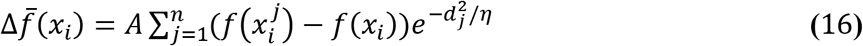

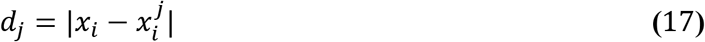

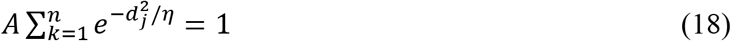

Where 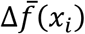 is the change in concentration of metabolite in grid space *x_i_* over a time step, *A* is the normalization constant to be calculated by solving the third equation within the entire neighborhood to ensure conservation of mass within the neighborhood, *d_j_* is the distance between grid space *x_i_* and its grid neighbor 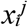 and *η* is the diffusivity control of the system over the time step. As *η* → 0 diffusion halts and as *η* → ∞ diffusion becomes instantaneous. In this study, 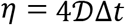 as in the original Fick’s Law.

Diffusion is calculated using two steps, one forward and one reversing the order of gridspace calculation, to mitigate the effect of order on estimating the concentration gradient (Figure S7). Excess ammonium is secreted into the environment using a membrane diffusion coefficient. Cells are allowed to uptake any metabolite/nutrient in YBC-II medium; the only extracellular products allowed in simulations are small molecules, such as CO_2_ and NH_4_^+^ which diffuse through the membrane, as well as compounds that have experimental evidence of transporters from proteomic analysis or transcriptomic analysis (estimated using membrane diffusion outwardly and free diffusion for gases or active transporters for ions/molecules inwardly) in Table S7 (67, 68).

The Ocean Agents also manage diffusion of metabolites from marine sinks and through the gas-liquid surface interface with the atmosphere. This is done assuming equilibrium concentrations of dissolved gases defined by Henry’s Law and mono-directional slab diffusion for CO_2_, O_2_, and N_2_. Table S6 lists the free diffusivities of compounds and Table S8 lists the Henry’s Constants for atmospheric compounds, and Figure S7 demonstrates the movement of diffusive molecules through the simulation.

Furthermore, light diffusion to cells is defined as a function of their y coordinate according to the equation:

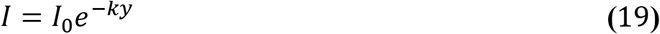

Where *I* is light intensity, k is the extinction coefficient of light in seawater, and y is the depth below the surface of the individual cell.

#### Filaments

Filament Agents are responsible for organizing cells, managing movement, splitting to promote diazotroph development, and defining cell type after division. Random walk movement (to simulate the lack of control cells have over lateral motion) is simulated by generating a random direction that has an empty grid space for every cell in the filament. Cells move within a user defined interval of time or if their growth is impeded by another filament, in which case growth is halted until the cells move away from each other. The filament forces splitting into two separate filaments when nitrogen is limiting growth and neither filament end is undergoing diazotroph development (meaning that another diazocyte is required). Filament Agents decide the next cell type using this inequality:

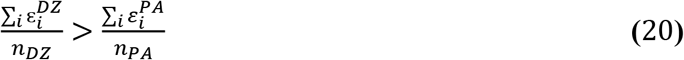

Where *ε* is the Pareto Efficiency of the given cell type and *n* is the quantity of that cell type in the filament. The Pareto Efficiency is quantified as the sum of the objective fluxes divided by their Pareto Optimum (from experimental results) divided by the number of objectives.

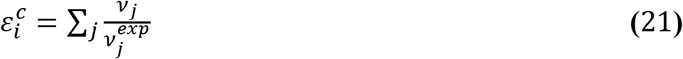

If inequality (20) is satisfied, the cell prioritizes diazotroph development, otherwise it prioritizes photoautotroph development. If a diazotroph region is currently under development, the filament adds another cell to that region. If there is no diazotroph under development, or if the C:N ratio becomes higher than physiological bounds, the filament splits to expose a region where diazotroph development may begin. A photoautotroph can be placed at any open site. Since there are two ends on every filament, up to two of these decisions are being made during each simulation time step. After filament splitting, if the split results in a homogenous region of either diazotrophs or photoautotrophs, the missing cell type is preferred. Filaments split in the middle of the longest region of homogenous cells and are prevented from splitting to result in a single cell, meaning that the shortest possible resulting splits are two cells in length. Cell division completes within one time step when metabolites and biomass are equally divided between parent and daughter cell and the filament updates to contain the cell at its end. This decision is a memoryless process conducted each time step. This means that cell division is completely metabolically motivated (which is affected, in turn, by diffusion and physiological processes).

### Parameter Estimation

As described previously, to improve the accuracy of simulations, the model was fit to experimental data for cells grown in 100 μE light in YBC-II medium. Maintenance energy, in the form of the ATP hydrolysis reaction, is the main parameter that is adjusted in FBA formulations (3, 69–72) to match simulations with growth rate. Since maintenance energy at 100 μE was higher than the energetic capacity of the model for growth at 50 μE, a linear correlation was interpolated from experiments at 100 μE and 80 μE with ATP maintenance flux fit to both cases for each cell type:

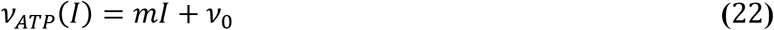

Where *m* and *ν*_0_ are calculated using the point-slope equation for a linear equation:

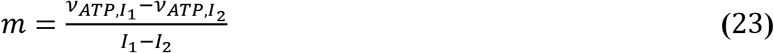

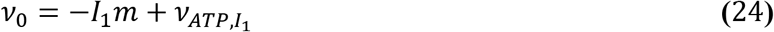

Where *I_1_* is 100 μE and *I_2_* is 80 μE. The estimated values of the linear equation are recorded below in Table 1. If the model is unable to satisfy its maintenance demand (through any metabolic process, including catabolizing its own biomass), the cell dies. *L_0_* is the energy required in zero light to maintain the cell without active metabolism.

**Table 1:** ATP maintenance flux requirements estimated as a function of light intensity for Pareto Fitting.

| Cell Type | Maintenance Energy, $v_{ATP}$ | | $m$<br>$\left(\frac{\text{mmol}}{\text{g DW h } \mu\text{E}}\right)$ | $v_0$<br>$\left(\frac{\text{mmol}}{\text{g DW h}}\right)$ | $L_0$ ($\mu\text{E}$) |
| --- | --- | --- | --- | --- | --- |
| | 80 $\mu\text{E}$<br>$\left(\frac{\text{mmol}}{\text{g DW h}}\right)$ | 100 $\mu\text{E}$<br>$\left(\frac{\text{mmol}}{\text{g DW h}}\right)$ | | | |
| Photoautotroph | 34.3 | 53.3 | 0.952 | -16.7 | 16.9 |
| Diazotroph | 62.3 | 82.0 | 0.987 | -41.9 | 44.0 |

### Multi-Objective Optimization

Unlike typical formulations of flux balance analysis (14, 71–76), which use a single objective function to predict fluxes, our model uses multi-objective optimization to more accurately approximate the true objectives of the cell: to optimize biomass while also producing the metabolite they exchange between cell types. Implementation of multi-objective optimization is more complex and computationally intense than single objective optimization therefore, to minimize computational effort, Pareto Fronts were generated *a priori* by iteratively increasing ATP maintenance flux and using every permutation of objective weights to fit to a dominant front (see *SI Methods* and Figure S8 for more complete details). For each point along the Pareto Front, Euclidean distance was used to determine the relative weight of each objective function, which was then used to generate a single, scalarized reaction. Each cell in the simulation calculates its scalar objective function separately during each time step based on its internal constitution and requirements.

### Implementation of Mutable Objective Functions

Previous studies have used static objective functions, where production is consistent during every phase of growth. However, organisms accumulate and digest metabolites during growth and development. To reflect this, we inserted a “mutable” objective function where relative preferences of storage compounds and biomass production can be tailored by the agent based on cell biomass. The scalarized objective equation was thus broken into two main components: storage compounds (cyanophycin modeled as β-aspartyl arginine and glycogen modeled as maltose) and biomass (lipids, proteins, DNA, RNA, chlorophyll, phycoerythrin, etc.). We assumed that biomass remained relatively stable throughout the day while the amount of storage compound was allowed to vary. The scalar weights, or production priorities, were manipulated assuming cells do not grow beyond twice their average cell without dividing: lower biomass prioritizes growth and higher biomass prioritizes vegetative storage compound production. Mathematically, this is modeled such that the scalar objective equation’s biomass coefficient was inversely adjusted by cumulative probability of a cell’s biomass in the distribution. The normal distribution was formulated assuming cubic 10 μm cells with density of water (77) as the average mass and a narrow distribution with a standard deviation of 0.433 times the mean size. This value was chosen to promote switch-like bistable behavior between cell phenotypes: either cells are biomass driven (exponential) or they are metabolite driven with combinations of probabilities in between. This is because a single sample of a cell from a distribution of cells would have a probability of 99% to fall between 0 and twice the mean size. The final distribution is:

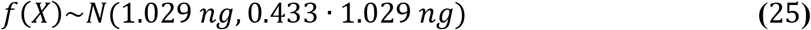

Calculation of new objective coefficients was done by first finding the cumulative probability (*z*) of another randomly selected cell’s non-metabolite biomass being less than or equal to the objective cell’s biomass at each time point for each cell:

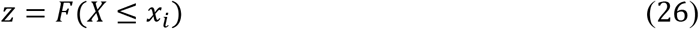

This is used to adjust the average, experimentally matched objective coefficient 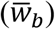 for biomass by multiplying that coefficient by the probability of the cell being larger than that size, a value that represents the probabilistic expansion space (*ϵ*) of the cell:

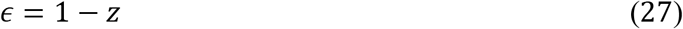

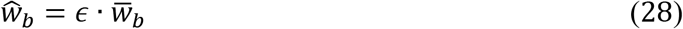

Major metabolite coefficients for the scalarized objective equation were also adjusted using this probability, increasing as the cell’s size increased:

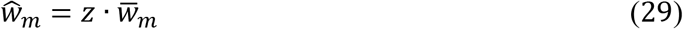

Finally, the coefficients are normalized such that:

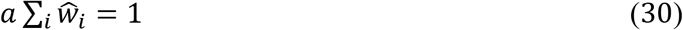

Or:

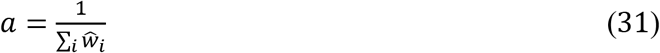

Which yields final objective coefficients of:

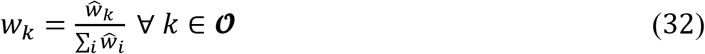

Where 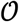 is the set of all objective metabolites in the original scalar equation.

Performance evaluation of the mutable objective function, validation of the mutable objective function versus the static version, and justification of non-metabolite biomass as the independent objective are provided in *SI Methods* and Figure S9.

## ABBREVIATIONS

6PG: 6-phospho-D-gluconate
6PGDL: 6-phosph-D-glucono-1,5-lactone
ABM: Agent-Based Modeling
AcCoA: Acetyl-CoA
AKG/αKG: α-ketoglutarate/2-oxoglutarate
ALA: L-alanine
βASP-ARG: β-aspartyl arginine
βG6P: β-glucose-6-phosphate
CBB: Calvin-Benson-Bassham Cycle
CDeg: Cyanophycin Degradation to Amino Acids
cEFMA: community Elementary Flux Mode Analysis
CIT: Citrate
COBRA: Constraint-Based Reconstruction and Analysis Toolbox
CobraPy: Constraint-Based Reconstruction and Analysis Toolbox for Python
COMETS: Computation of Microbial Ecosystems in Time and Space
CSV: Comma Separated Value
cph: Cyanophycin
CSyn: Cyanophycin Synthesis from Amino Acids
dFBA: dynamic Flux Balance Analysis
DHAP: Dihydroxyacetone phosphate
DON: Dissolved Organic Nitrogen
E4P: Erythrose-4-phosphate
EF: Efflux
EX: Export
F6P: Fructose-6-phosphate
FBA: Flux Balance Analysis
FDP: Fructose 1,6-diphosphate
FOR: Formate
FUM: Fumarate
FVA: Flux Variability Analysis
G6P: Glucose-6-phosphate
GAP: Glyceraldehyde 3-phosphate
GDeg: Glycogen degradation
GLX: Glyoxylate (flux maps) or Glyoxylate and dicarboxylate metabolism (Figure 4)
GLY: Glycine (flux maps) or Glycogen/Gluconeogenesis (Figure 4)
gly: Glycogen
GLYR: Glycerate
GOL: Glycerol
GP: 3-phosphoglycerate
GSyn: Glycogen synthesis
ICIT: Isocitrate
IN: Influx
JSON: JavaScript Object Notation
jyCOBRA: java-python integrated COBRA
KEGG: Kyoto Encyclopedia of Genes and Genomes
LIP: Lipid metabolism
MAL: Malate
MSM: Multiscale Modeling
OAA: Oxaloacetate
PEP: Phosphoenolpyruvate
PGOL: Phosphoglycolate
Pi: Inorganic phosphate
PPP: Oxidative Pentose Phosphate Pathway
PRO: Protein synthesis
PYR: Pyruvate
R5P: Ribose-5-phosphate
REF: Reflux
Ru5P: Ribulose-5-phosphate
RuBP: Ribulose 1,5-bisphosphate
S17P: Sedoheptulose 1,7-bisphosphate
S7P: Sedoheptulose 7-phosphate
SBML: Systems Biology Markup Language
SUCC: Succinate
SUCSAL: Succinic semialdehyde
SDA: Subsystem Distribution Analysis
TCA: Tricarboxylic acid cycle
TCP/IP: Transmission Control Protocol/Internet Protocol
X5P: Xylulose 5-phosphate

## DECLARATIONS

### Availability of data and materials

The ABM framework and datasets resulting/analyzed from simulations are available on the GitHub repository at https://github.com/boylelab/iTery101-ABM with username “boylelab” and password “diazo101”. We will make this public upon publication.

### Author contributions

JJG, BMSH and NRB designed the research. JJG performed the research. JJG and NRB analyzed the data. JJG, BMSH and NRB wrote the manuscript.

## Acknowledgements

Not applicable.

