## Supplemental Data for "MultIscale MultiObjective Systems Analysis (MIMOSA): an advanced metabolic modeling framework for complex systems"

### I. Model Formulation

I.A Genome-Scale Model

Investigations suggest that filamentous, diazotrophic cyanobacteria store nitrogen as cyanophycin and passes it to photoautotrophic cells as the cyanophycin derivative β-aspartyl arginine (2). We modeled this phenomenon by adding $\beta$-aspartyl-arginine anabolizing and catabolizing pathways to the model. These pathways (and their genetic evidence) are listed in Table S2. The requirement of cyanophycin catabolism also required aspartate catabolism. Genomic evidence suggested that arginine is deaminated to ornithine. While no specific ornithine deamination pathways are suggested by genome annotation, amino acid deaminases tend to be promiscuous. Therefore, the arginine catabolism was assumed to progress through succinic-semialdehyde (4-oxobutanoate) and into the TCA cycle, yielding TCA cycle metabolites and glutamate.

Export and transport reactions for $\beta$-aspartyl arginine were also added to add mass balance closure to the model (similar to the biomass transport and export equations). Concurrently, cyanophycin and glycogen contributions were removed from the biomass equations, effectively decoupling primary metabolites from the biomass equation and allowing them to function separately within the optimization algorithm.

Additional energetic limitations are placed on the models to resemble realistic conditions: light is constrained to 100 $\mu$E to reflect laboratory results (3). Glycogen and cyanophycin uptake by diazotrophic and photoautotrophic cells, respectively, was estimated using composition measurements and mass balances. ATP maintenance bounds were determined via the correction equation detailed in the “Pareto Front Generation” section and the linearized light dependent Equation 27 with values recorded in Table 1.

I.B Pareto Front Generation

A single, discrete solution does not exist for these bi-objective formulations. This is addressed through an empirical, *a posteriori* method for matching theoretically generated cell production to measured results. First, a Pareto Front is generated using multi-objective flux balance analysis adapted from Pareto Front generation of previous optimization problems (4, 5). Each objective was given a weighting coefficient such that the sum of the coefficients summed to one. Every combination of coefficients satisfying this constraint are applied to the scalarized objective to generate the Pareto Front. Then, a scalarized objective function corresponding to the sum of every metabolite of every participating reaction with their original coefficient multiplied by the scalar weight is added to the model. This is of the form:

$\mathbf{max}\{\sum_{\boldsymbol{i}} a_{i}\nu_{i}\}$	[S1]

subject to $\sum_{\boldsymbol{i}} a_{i}=1$	[S2]

$\boldsymbol{S}\cdot\boldsymbol{v}=\boldsymbol{0}$	[S3]

$\boldsymbol{v}_{\boldsymbol{ex}}=\mathcal{E}$	[S4]

Where $a_{i}$ references the scalar coefficient for reaction flux $\nu_{i}$. ***S*** and $\boldsymbol{\nu}$ are the stoichiometric and flux matrices. $\boldsymbol{\nu}_{\boldsymbol{ex}}$ is the set of extracellular fluxes and $\mathcal{E}$ is the set of experimentally measured consumption rates.

The Pareto Front is plotted with each objective corresponding to an axis. Dominated points (points for which there can be a direct improvement in each objective) are discarded. This application was for two bi-objective problems, but the formulation is applicable to higher order formulations as well.

Once the Pareto Front is generated, it is fit to experimental data. This is done by using a constraining equation – in this case, ATP hydrolysis – to summarize futile cycles and other non-metabolism factors that might reduce metabolic efficiency in experimental conditions. ATP hydrolysis has been used in prior genome-scale reconstructions for these purposes (6) and is iteratively increased until the Euclidean Distance between the theoretical Pareto Front and the actual, average experimental data for a given condition is minimized. The scalarized objective function accounting for the closest theoretical coordinate is selected and used as a scalar function for multi-objective optimization. The goal values and constraints for each cell type and each light regime are listed in Table S3 and are taken from previous work (5) in addition to this.

These calculations are conducted using CobraPy (7) for Linear Optimization and SBML processing and NumPy (www.numpy.org) for non-routine mathematical operations (like square roots and matrix math). Pandas (pandas.pydata.org) is used to store solutions to a CSV table as they are created. *In situ* and experimental data are read via CSV and are used to update the bounds and objective functions for the model. JPServe creates a locally hosted TCP/IP connection to pass JSON encoded arrays – containing the scalarized model, reaction fluxes, and cell identification information (position, filament size, time step, and type) – between Python (for Pareto and FBA calculations) and Java (for the Agent Based Model) (John Huang, https://github.com/johnhuang-cn/jpserve).

I.C **Shifting Scalar Objectives**

To investigate the mutability of the scalar objective equation, and the effects of intra- and inter-cellular conditions on cell objectives, an algorithm to shift the objective function along the Pareto Curve is implemented based on the biomass of the cells (assuming that structural biomass is relatively intransigent in these conditions). That means that, as biomass accumulates, a cell will behave more stationarily, i.e., it will prioritize secondary metabolites. This is adjusted assuming a normal distribution in cell sizes:

$f\left( X \right)\sim N(\mu,\sigma)$	[S5]

where $\mu$ is the average biomass (assuming cubic shape and density near water) or metabolite concentration (from experimental data) and $\sigma$ is the standard deviation. As the standard deviation decreases, the model is less tolerant of deviations from mean values and will more readily adjust the scalar weights. As it increases, it will be more tolerant of deviations from mean values and will cause smaller perturbations to scalar weights. For this study, the standard deviation was assumed to be 0.433 so that 95% of cells would fall within a factor of seven from minimum to maximum and a mean of 1.029 $\times{10}^{-9}$ g calculated using an assumed cubic shape with dimensions of 10 $\mu m$ per side and a density of seawater (8).

The correction occurs by using the cumulative distribution function for the normal distribution. Given a Pareto matching algorithm resulting set of weights, the weights are shifted (inversely) as:

$w_{1}=a\hat{w}_{1}$	[S6]

$\hat{w}_{1}=\bar{w}_{1}\left[ 1-F\left( X_{1}=x_{1} \right) \right]$	[S7]

$w_{j\neq1}=w_{j\neq1}F\left( X_{j\neq1}=x_{j\neq1} \right)$	[S8]

$a\sum_{i} w_{i}=1$	[S9]

Where $w_{1}$ is the new, shifted weight, $\hat{w}_{1}$is the non-normalized and shifted weight, and $\bar{w}_{1}$ is the Pareto matched weight. X is the variable and x is the cell’s quantity. Meaning, in each case, that as the objective increases, it responds by producing the corresponding objective: as biomass increases, it expends more resources on biomass.

I.D System Variables

The model initializes a tunable number of identical filaments containing a tunable number of diazotrophs and photoautotrophs. These are populated at random coordinates on a two-dimensional wrapping grid (meaning that x^MAX^ = x^0^) signifying width (X) and depth (Y) coordinates. Thus, each coordinate represents one grid cell and therefore one grid cell length. Furthermore, the names of the transactional metabolites (diazotroph → photoautotroph and photoautotroph → diazotroph) as well as the name of the biomass equation are given by the user when the simulation is initialized. The time-step length, governing diffusion coefficient (for determining neighborhood bounds), and random-walk move times to simulate drifts due to current and inter-filamental contact. Finally, model and data file path information is included. The full set of user defined parameters and their default values is summarized in Table S4.

I.E **Routine Metabolic Optimizations**

Once the model is scalarized according to an experimental condition, it is optimized according to this objective in conjunction with constraints from concentration information from the ocean and filament sharing cell agents in the model. Metabolite concentrations in the cell agents are interpreted into fluxes and vice versa using:

$\nu=\frac{\Delta N}{X \Delta t}$	[S10]

Where $\nu$ is flux in mmol (g DW h)^-1^, ΔN is the change in the number of millimoles over the time step, X is cell biomass in g, and $\Delta$t is the time step duration in h.

The available environmental concentrations therefore form constraints on uptake. These updated models are solved according to these constraints and to the scalarized objective function, resulting in a series of flux solutions. The internal metabolic fluxes are stored to a CSV file. The external fluxes are then interpreted into consumed and produced metabolite concentrations over the period and are used to update the both intracellular and oceanic concentrations. Objective flux is the consolidated scalarized flux of all objective metabolites for a cell type (biomass and cyanophycin for diazotrophs/biomass and glycogen for photoautotrophs) and higher objective fluxes represent higher metabolic production. These are reinterpreted into discrete fluxes using the weighted objective equation.

Optimization of metabolism via the scalarized objective function results in three major cellular outcomes: anabolism, primary metabolite production, and catabolism. The cell prioritizes regimes in that order: anabolism, if unavailable, results in using the same nutrients for simply production and, when that is unavailable, catabolizes biomass for maintenance energy. The catabolic model was generating by iterating through necessary reactions for metabolic backflow from biomass to the primary metabolite and ATP and then creating a separate set of constraints for this situation.

Overall major metabolite passage is visualized in Fig. S7. Nitrogen gas is reduced and incorporated into amino acids, ultimately leading to the cyanophycin precursor $\beta$-aspartyl arginine. Carbon dioxide is reduced via the Calvin Cycle and incorporated into long sugars, modeled as the disaccharide maltose. Each unique cell type can pass its metabolite to the other cell type. However, each cell type also maintains the machinery to metabolize the reduced intermediates between the oxidized gases and their ultimate polymers (depicted by the red lines) as sources for their own objective metabolites. This represents a potential futile cycle where cyanophycin can be used primarily as a carbon source and not as a nitrogen source.

Therefore, to mitigate futile cycling of resources (one primary metabolite being burned excessively during vegetation to create the other primary metabolite), the maximum uptake of cyanophycin is modified as:

$\nu_{cph EX}\leq-\frac{N_{cph}}{X\Delta t}F$	[S11]

Where F is a tunable parameter for fraction of maximum uptake is allowable for uptake by a cell for catabolism over a timestep. F was proposed to be 0.08 and was corroborated through both sensitivity analysis and through experimental estimation from the similar organism *Anabaena*, which also exhibits specialized cell production of cyanophycin granules. This estimation was done using the ratio of heterocyst/vegetative exchange coefficients (0.022 s^-1^) and vegetative/vegetative exchange coefficients (0.29 s^-1^) to predict how cyanophycin accumulation impairs its dispersal during filamental development (2).

I.F State Recapture

Minimizing computational burden is a major concern for large-scale metabolic models. To enable computational efficiency while preserving high resolution of the system, a “state recapture” algorithm was developed. The state of the community at every time step is recorded in a CSV file. The program reads in the data and assigns an identity, location, nutrient profile, and filament to each cell. The user can then specify which community state with which to restart the simulation, and the user has latitude to adjust input parameters to either simulate a sudden system change (introduction of a nutrient or a cataclysm) or to increase the resolution of the time step. This latter effect allows the user to “zoom in” on a behavior or growth region of interest. Furthermore, the user can compare several different assumptions at once: this state recapture can be designed around a specific growth region and a parameter – for example, the diffusion constant to determine neighborhood size – to perform sensitivity analysis.

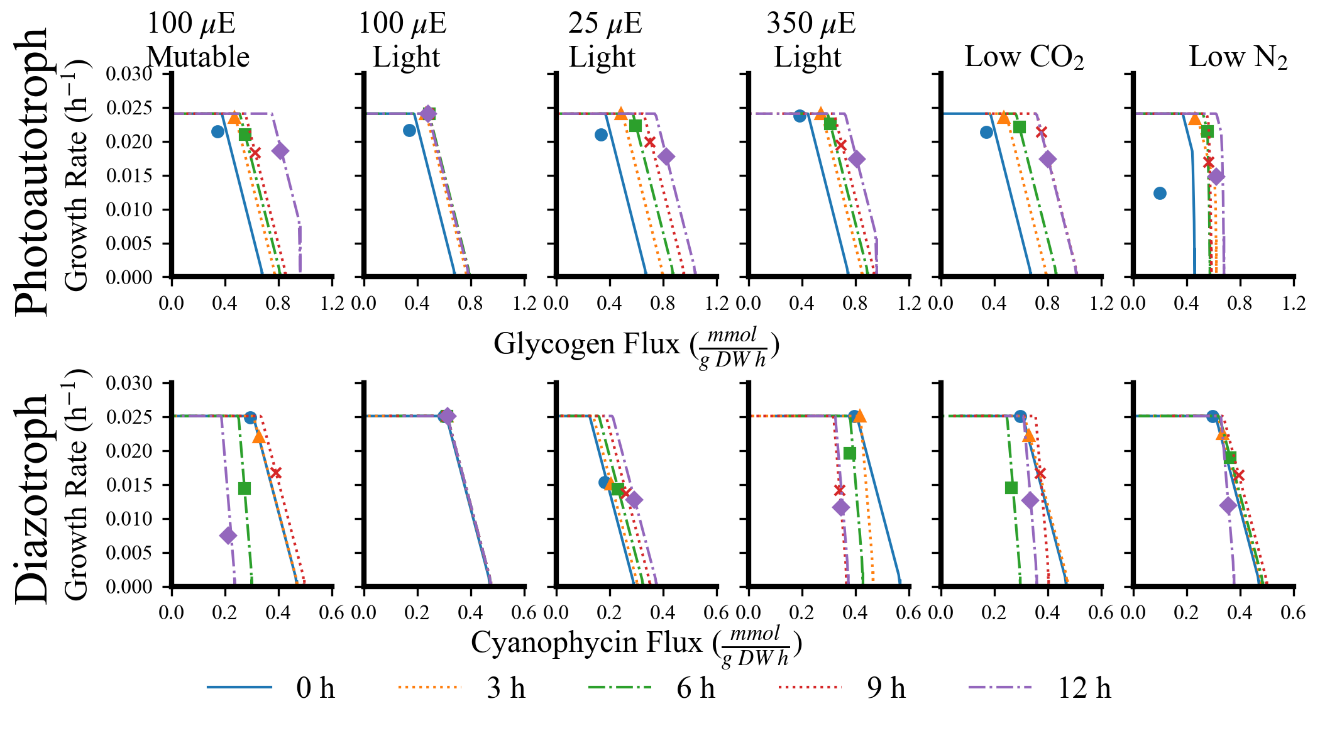

Figure S1. Derivation of scalar functions from all possible biobjective combinations as visualized by a Pareto Front in each simulation context. Progressing from left to right are different simulations conducted in atmospheric conditions, YBC-II media, under 100 $\boldsymbol{\mu}$E light, and with the mutable objective equation unless otherwise stated. Low CO_2_ and low N_2_ describe 200 ppm (versus 400 ppm) CO_2_ and 39.5% (as opposed to 79%) N_2_. Differently colored lines are different points during the light cycle for each simulation, each line describing a 3-hour increment. Points describe the selected objective equation at each time point. Top row indicates photoautrophic cell responses, bottom row indicates diazotrophic cell responses.

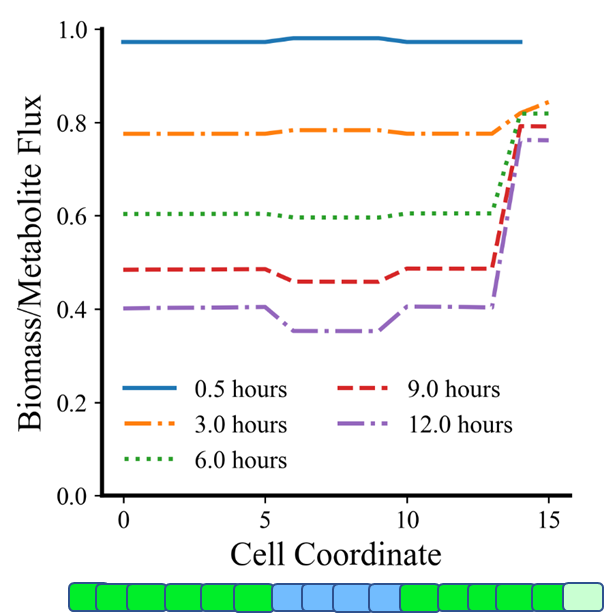

Figure S2. Evolution of Metabolic Prioritization within the Filament during the Day. Each color represents a later time point in the simulation. The y-axis corresponds to the normalized ratio of biomass scalarized weight to metabolite scalarized weight. 1.0 represents the biomass to metabolite ratio consistent with training data. Cell coordinates progress left to right along the filament, as visualized by the model filament beneath. Clear differences can be seen between photoautotrophic (peripheral) and diazotrophic (interior) cells. Division occurs at ends and new cells prioritize growth to metabolite development, visualized by the lighter colored green cell at the right end of the model filament.

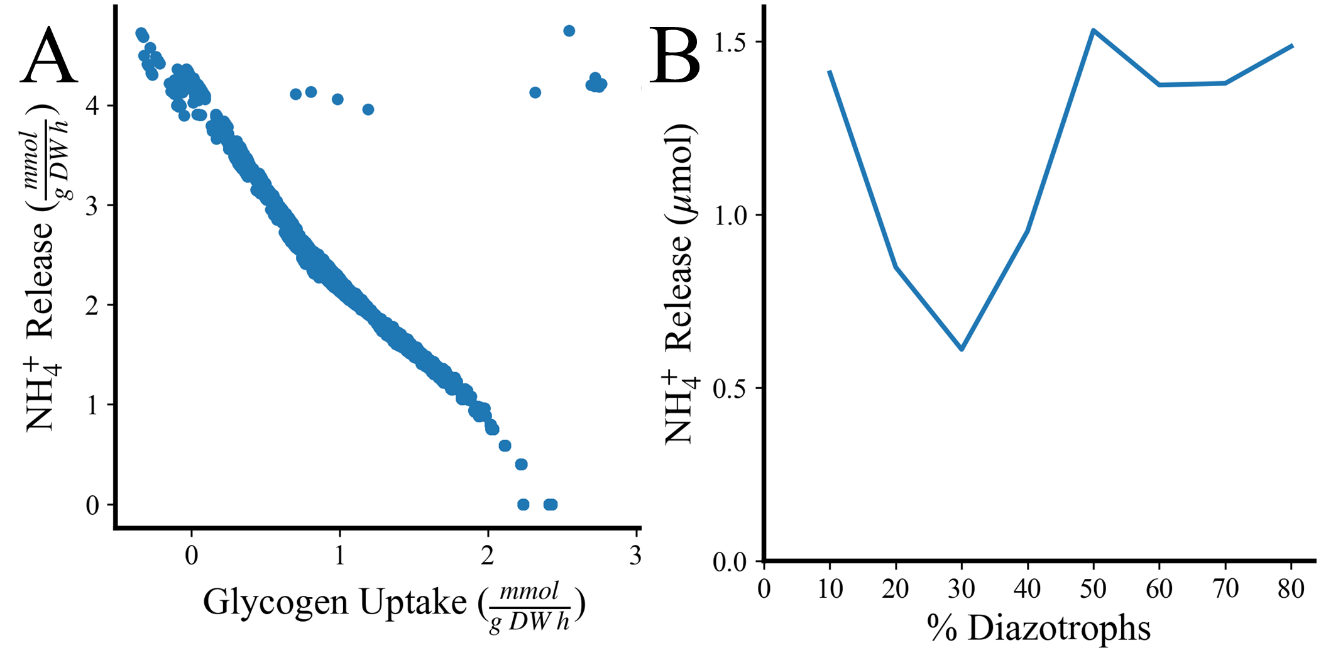

Figure S3. Ammonium Release as a Function of Metabolic and Population Imbalances. A) Ammonium flux out of the cell as a function of glycogen uptake flux. B) Total ammonium released into the media as a function of the percentage of each filament that is diazotrophic cells.

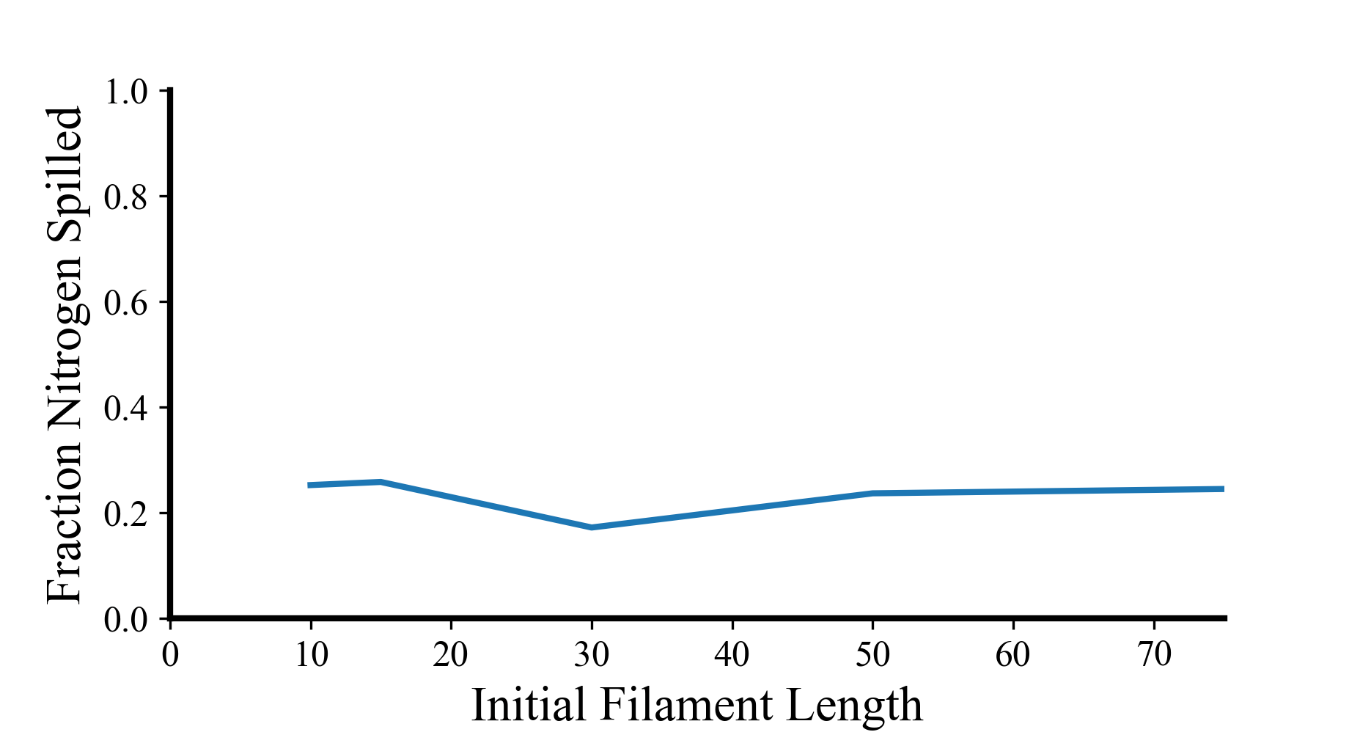

Figure S4. Fraction Fixed Nitrogen Spilled as a Function of Initial Filament Length. Percent of nitrogen released versus uptaken, using the computationally analogous method to the experimental method presented in (1). Nitrogen release varies little with different initial filament lengths, but remains between ~0.2 to ~0.3 of the total nitrogen uptaken. Simulation was conducted with constraints illustrated above and with ratios of 7:3 photoautotrophs: diazotrophs.

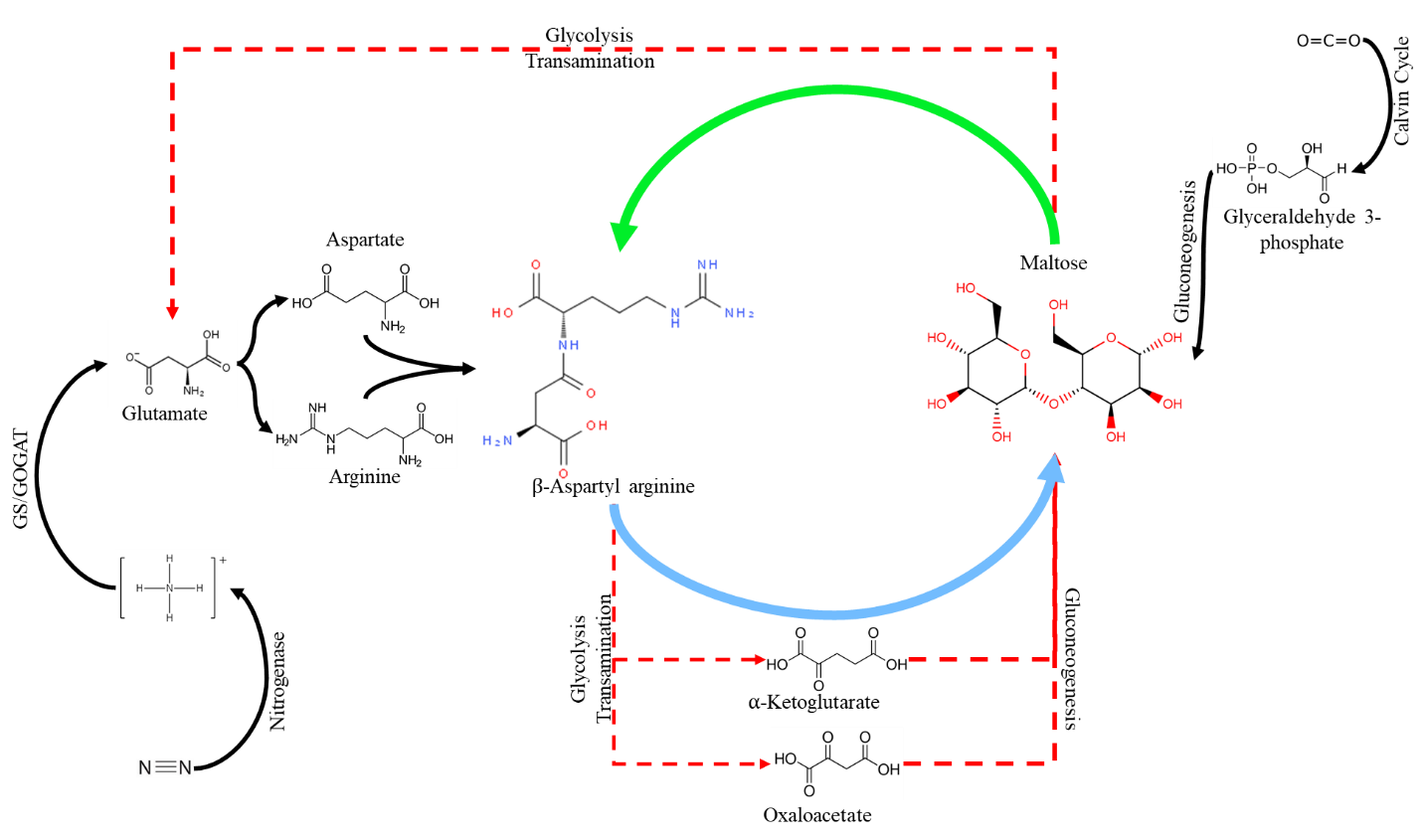

Figure S5. Metabolite Cycling of Major Nitrogen and Carbon Source Polymers in *T. erythraeum*. Black lines represent common metabolic processes between molecules. The thick green line and thick blue line represent trade from the photoautotroph and diazotroph, respectively. Dashed red lines represent potentials for futile cycling of transactional metabolites.

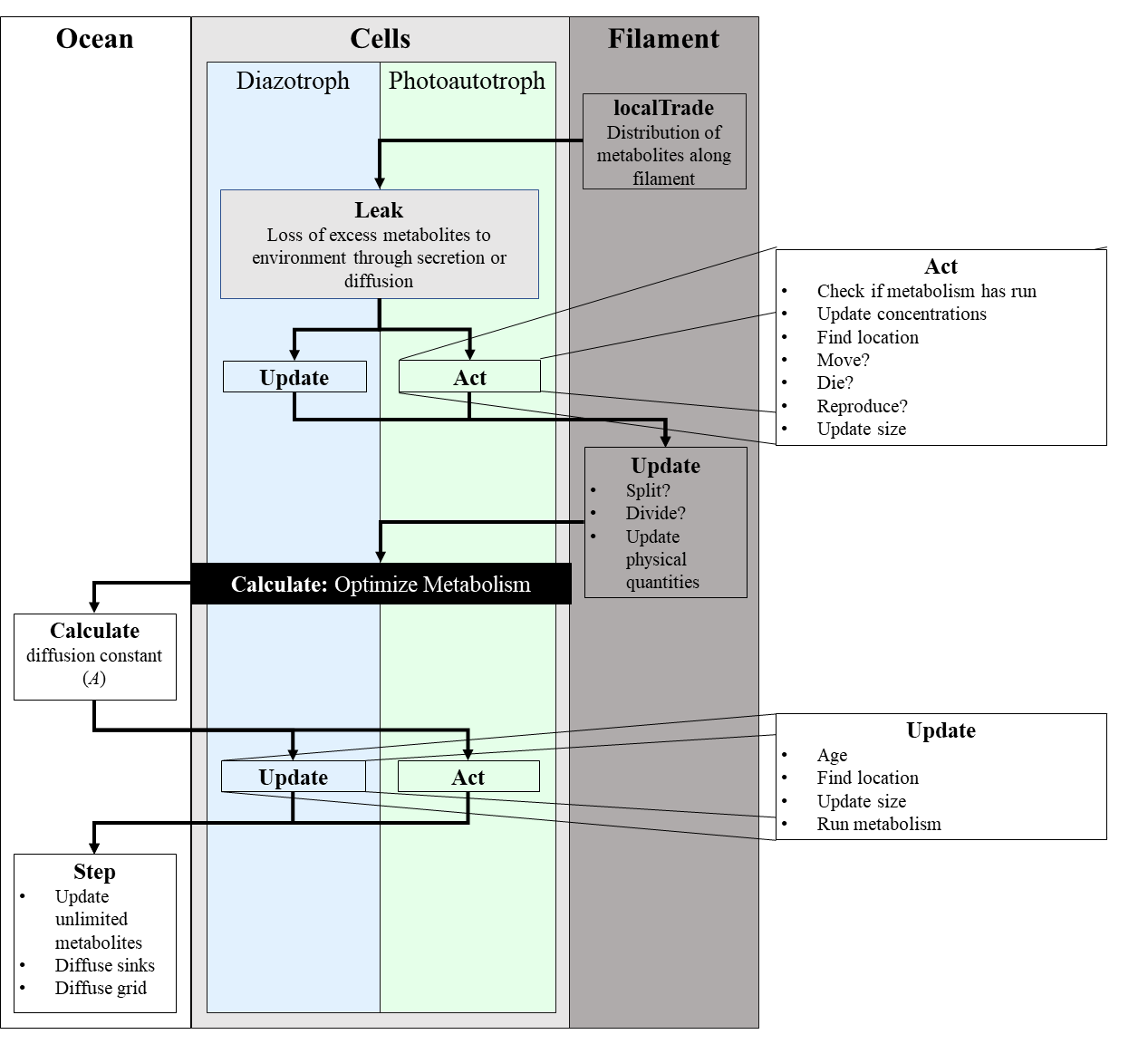

Figure S6. General Flow Chart of agent based decision making as the simulation progresses during one time step. From left to right: the white box are ocean agents which govern environmental diffusion and delivery of metabolites to near cells. The light gray box containing the blue and green boxes are cell agents. All cell agents leak metabolites, update physical quantities and processes, and act in the same way, except that they employ different metabolic models. The black box is a special class that handles all interactions with the Python modules that optimize the model. Moreover, photoautotrophs and diazotrophs are on opposite update/act schedules so that one type is always reacting to the other. Finally, the dark gray box indicates filament agents which govern dividing rules, cell organization, and intercellular, intrafilamental diffusion. Offset boxes indicate the specific tasks conducted by cells during update and act steps.

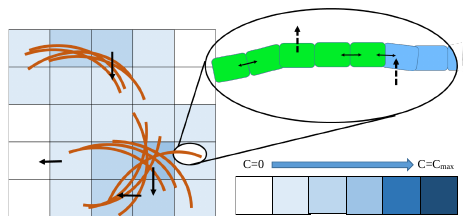

Figure S7. Scheme for Modeled Diffusion. The medium is modeled as a grid space with individual space dimensions of 10 grid cells $\boldsymbol{\times}$ 10 grid cells (100 $\boldsymbol{\mu}$m $\boldsymbol{\times}$ 100 $\boldsymbol{\mu}$m) and a total grid space of dimensions 2500 $\boldsymbol{\mu}$m $\boldsymbol{\times}$ 2500 $\boldsymbol{\mu}$m (250 grid cells $\boldsymbol{\times}$ 250 grid cells) and are modeled as a freely diffusive, dilute environment. This behavior exists within filaments; nutrients are able to freely diffuse between cells. Free diffusion between dilute environments is indicated with a solid black line. Green cells are photoautotrophs and blue cells are diazotrophs. Darker blue indicates higher concentration. Dotted lines indicate membrane mediated diffusion, demonstrating that permeation out of and into the cell occurs orders of magnitude more slowly unless there is evidence of an active transporter.

Figure S8. Algorithm for determination of an experiment-fitted Pareto Front, creation of a scalarized objective function, and objective adjustment for mutable function scaling. A) Generation of the average objective function. First, the theoretical Pareto Front is generated. Then, through adjustment of a corrective reaction (usually ATP hydrolysis), the Pareto Front is matched to the experimental data points. Finally, the points are weighted and interpreted into specific weights for the optimization function (black line). This is shown in two-dimensions for *T. erythraeum* but is theoretically viable for many dimensions of non-dominated Pareto Fronts. B) Illustrates the general scheme of determining the objective function. First, the Pareto Curve is generated from the Genome Scale Model and the given objectives. Then, the biomass maintenance flux is increased until the Euclidean distance between experimental points and the curve is minimized. Then, the corresponding point on the curve is used to generate the scalarized objective function. C) Demonstrates how the Pareto Curve might change based on an environmental change (nitrogen limitation) and how the objective function most similar to the empirical data set may change as well. Instead of generating a new Pareto Curve for every circumstance, a normal distribution-based weighting function corrects for these scenarios instead.

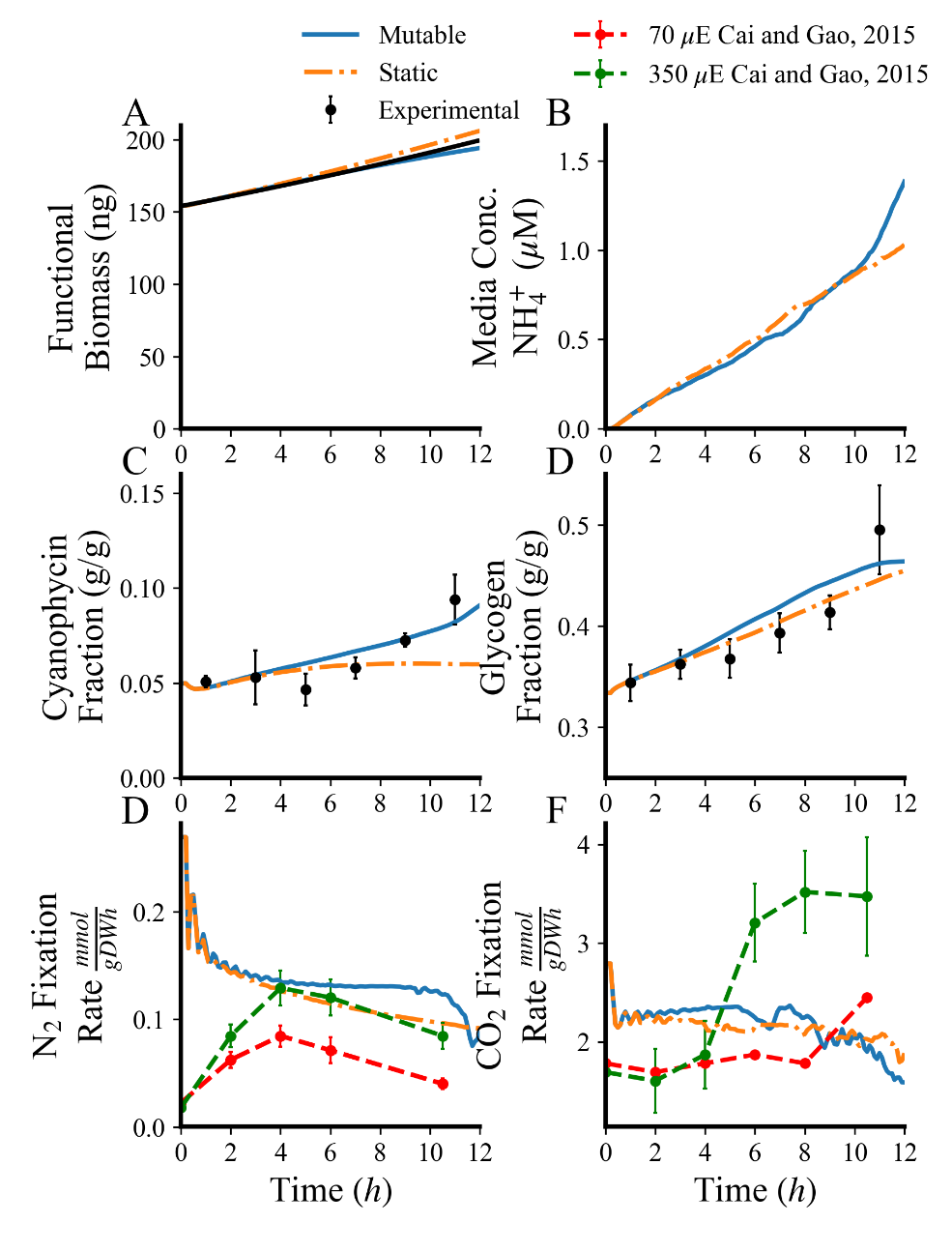

Figure S9. Contrasting Performances of Mutable and Static Objective Functions. Blue lines indicate weighted (mutable) objective functions that respond to cell composition and orange lines indicate unweighted (static) objective equations. The simulations were run in conditions modeled after empirical data collection (YBC-II media, atmospheric nitrogen and carbon, 100 $\boldsymbol{\mu}$E light). A) Non-metabolite growth rates (lumped biomass constituents that are not cyanophycin or glycogen, i.e., proteins, lipids, RNA, DNA, etc.). The unweighted biomass demonstrates a higher non-metabolite growth rate than the weighted simulation. B) NH_4_^+^ leakage in the mutable objective simulation follows the same pattern during the first part of the simulation, but shows exponential tendencies at the end, while the static objective remains linear. C) Cyanophycin biomass fraction in mutable and static simulations. The mutable objective equation simulation is able to recover effectively from initial low biomass fraction of cyanophycin while the static objective tails off logarithmically and cannot reach appropriate fractions of cyanophycin. D) Glycogen accumulation is higher across the entire time period in the mutable objective case than in the static one and shows non-linear accumulation trends later in the growth cycle that mirror experimental evidence. E) Nitrogenase rate does not display the same initial delay as in experimental evidence, probably indicating a regulatory element not reflected in this model, but does not tail off in the simulation like in nature. The simulation with the mutable objective function fixes nitrogen at a consistently higher rate. F) Carbon dioxide simultaneously activates early in the light cycle for both simulations and remains fairly constant in both mutable and static objectives. This mirrors lower late conditions, implying that the model has a floor of carbon fixation activity that is a primary and relatively static objective.

Table S1. Reactions/Enzymes Added to Genome Scale Model

| **Proposed function** | **E.C. number** | **Gene** | **Annotated function** | **Source organism** |
| --- | --- | --- | --- | --- |
| Cyanophycin synthetase (aspartate adding) | 3.4.15.6 | Tery_1964 | Cyanophycin synthetase | *Trichodesmium erythraeum* |
| Cyanophycin synthetase (arginine adding) | 3.4.15.6 | Tery_1964 | Cyanophycin synthetase | *Trichodesmium erythraeum* |
| Cyanophycin synthetase (initiation) | 3.4.15.6 | Tery_1964 | Cyanophycin synthetase | *Trichodesmium erythraeum* |
| Cyanophycinase (aspartate removing) | 3.4.15.6 | Tery_1964 | Cyanophycinase | *Trichodesmium erythraeum* |
| Cyanophycinase (arginine removing) | 3.4.15.6 | Tery_1964 | Cyanophycinase | *Trichodesmium erythraeum* |
| Cyanophycinase ($\beta$-aspartyl arginine hydrolyzing) | 3.4.15.6 | Tery_1964 | Cyanophycinase | *Trichodesmium erythraeum* |
| L-Arginine carboxylyase | 4.1.1.19 | Tery_1142 or Tery_1276 | Arginine decarboxylase | *Trichodesmium erythraeum* |
| Agmatine amidinohydrolase (agmatinase) | 3.5.3.11 | Tery_3780 | Agmatinase | *Trichodesmium erythraeum* |
| Putrescine aminotransferase | 2.6.1.82 | Tery_2649 | Acetylornithine aminotransferase | *Amycolatopsis methanolica* |
| S-Adenosylmethioninamine putrescine 3-aminopropyltransferase | 2.5.1.16 | Tery _0722 or  Tery _0248 or Tery_0886 | Spermidine synthase | *Trichodesmium erythraeum* |
| Aminobutyraldehyde dehydrogenase | 1.2.1.19 | Tery_2599 | Aldehyde dehydrogenase | *Trichodesmium erythraeum* |
| 4-Aminobutyrate transaminase | 2.6.1.19 | Tery_2649 | Aspartate aminotransferase family protein | *Nostocales cyanobacterium* HT-58-2 |

Table S2. Goal Values for Model Training at 100 $\boldsymbol{\mu}$E and 80 $\boldsymbol{\mu}$E

| **Light Intensity (µE)** | **Cell Type** | **Growth Rate (h^-1^)** | **β-Aspartyl Arginine Flux (mmol (g DW h)^-1^)** | **Maltose Flux**  **(mmol (g DW h)^-1^)** |
| --- | --- | --- | --- | --- |
| 80 | Diazotroph | 0.0216 | 0.369 | -0.699 |
| 80 | Photoautotroph | 0.0216 | -0.0478 | 0.244 |
| 100 | Diazotroph | 0.0226 | 0.386 | -0.731 |
| 100 | Photoautotroph | 0.0226 | -0.05 | 0.255 |

Table S3. Flux Balance Analysis Constraints

| **Reaction** | **Lower Bound**  **(mmol (g DW h)^-1^)** | **Upper Bound**  **(mmol (g DW h)^-1^)** | **Source/Reason** |
| --- | --- | --- | --- |
| Bicarbonate uptake | -2.04 | 0 | YBC-II media (9) |
| Calcium uptake | -10.0 | 0 | YBC-II media (9) |
| Sulfate uptake | -25.0 | 0 | YBC-II media (9) |
| Sodium uptake | -0.411 | 0 | YBC-II media (9) |
| Magnesium uptake | -20 | 0 | YBC-II media (9) |
| Light uptake | -100 | -100 | Laboratory Data |
| CO_2_ uptake | -0.778 | 0 | Atmospheric conditions (10) |
| N_2_ uptake | -48.0 | 0 | Atmospheric conditions (10) |

Table S4. Cell Variables, Purposes, and Ranges

| **Variable** | **Purpose** | **Default Value or Range** |
| --- | --- | --- |
| **Static (Class) Variables** | | |
| AVE_WEIGHT | Average *T. erythraeum* cell weight | 1.029 ng (Assuming a cubic shape, a density near water, and a length of 10 $\mu$m (23)) |
| EXTINCTION_COEFFICIENT | Extinction coefficient for sunlight in saltwater | 2.625 x 10^-3^ m^-1^ (24) |
| SURFACE_LIGHT | Light at the surface of the ocean | 100 μE (3) |
| CELL_LENGTH | Average estimated cell length of *T. erythraeum* | 10 μm (25) |
| CONC_CPH_START_PER_G_DW | Starting concentration of cyanophycin within cell | 1.026 mmol (g DW)^-1^  0.33 g (g DW)^-1^ (This study) |
| CONC_GLY_START_PER_G_DW | Starting concentration of glycogen within cell | 0.172 mmol (g DW)^-1^  0.033 g (g DW)^-1^ (This study) |
| massDist | Mass distribution of cells | $F\left( X \right)\sim N(\mu,\sigma_{B})$ |
| MEMBRANE THICKNESS | Thickness of the lipid bilayer | 70 nm (17) |
| TIME_STEP | Duration of time step | 0.1 h |
| biomass | Name of biomass equation specified by user | “biomass” |
| CPH_MM | Molar mass of cyanophycin | 0.28929 g/mmol |
| GLY_MM | Molar mass of glycogen | 0.32529 g/mmol |
| BioF | Cyanophycin uptake restriction factor | 0.08 |
| **Individual Cell (Local) Variables** | | |
| myFilament | Filament of which the cell is a part | Inherited from progenitor cell |
| canDiazo | Boolean describing whether the cell is able to divide into a diazotroph | False |
| pools | Nutrient concentrations in cells | Initialized with ideal concentrations |

Table S5. Selected Permeabilities for Cellular Metabolites Through a Lipid Bilayer

| **Compound** | **Permeability (cm/s)** | **Modeled Diffusivity**  **(70 nm wall (17)) (**$\boldsymbol{\mu}$**m^2^ s^-1^)** |
| --- | --- | --- |
| NH_4_ | 4.80 $\times$ 10^-2^ (18) | 0.336 |
| O_2_ | 1.14 $\times$ 10^6^ (19) | 7.98 $\times$ 10^6^ |
| CO_2_ | 3.5 $\times$ 10^3^ (20) | 250 |
| Urea | 6.1 $\times$ 10^-7^ (21) | 4.3 $\times$ 10^-8^ |

Table S6. Diffusivities of Seawater Constituents and Metabolites

| **Compound** | **Diffusivity (**$\boldsymbol{\mu}$**m^2^ s^-1^)** |
| --- | --- |
| NH_4_^+^ | 2070 (11) |
| CO_2_ | 1880 (12) |
| O_2_ | 2000 (11) |
| N_2_ | 1880 (13) |
| CO | 2030 (13) |
| H_2_ | 4500 (13) |
| Cl^-^ | 1771 (14) |
| Na^+^ | 1212 (14) |
| SO_4_^2-^ | 803 (14) |
| HCO_3_^-^ | 1000 (14) |
| Ca^2+^ | 729 (14) |
| Mg^2+^ | 980 (15) |
| Mn^2+^ | 688 (15) |
| Fe^3+^ | 607 (15) |
| Fe^2+^ | 719 (15) |
| Co^2+^ | 699 (15) |
| K^+^ | 2070 (15) |
| Cu^2+^ | 733 (15) |
| NO_3_^-^ | 1700 (16) |

Table S7. Included Extracellular Transporters

| **Reaction ID** | **Reaction Name** |
| --- | --- |
| TR_arsi | Oxyanion-translocating ATPase/Arsenite efflux pump ARC3 |
| TR_bet | ABC-type dipeptide/oligopeptide/nickel transport: betaine |
| TR_biomass | Transport biomass |
| TR_biot | ABC-type dipeptide/oligopeptide/nickel transport: biotin |
| TR_ca2 | Calcium/proton antiporter |
| TR_co | CO transport via diffusion |
| TR_co2 | CO_2_ transport via diffusion |
| TR_cobalt2 | ABC-type cobalt transport |
| TR_cu2_abc | Copper transport via ABC system |
| TR_fe2 | Iron (II) transport via ABC system |
| TR_fe2_Sp | Iron (II) Transport System |
| TR_fe3 | Fe^3+^-BC transport |
| TR_h2 | Hydrogen Export |
| TR_h2o | H_2_Ot5 |
| TR_k_ACT | Potassium uptake Active |
| TR_k_SYM | Proton Antiport Potassium |
| TR_mg | Magnesium transport via ABC system |
| TR_mg_SP | Magnesium transport via MgtE |
| TR_mn2 | Manganese-ABC transport |
| TR_n2 | N2 transport |
| TR_na_h | Proton sodium antiport |
| TR_nh4 | Ammonium permease |
| TR_no2 | Nitrite transport in via proton symport |
| TR_hco3 | ABC-type bicarbonate transport |
| TR_Ca2Na | Calcium/Sodium antiporter |
| TR_pi_ABC | ABC-type phosphate |
| TR_no3 | Nitrate transport in via proton symport |
| TR_no3abc | Nitrate transport in via ABC system |
| TR_o2 | O_2_ transport via diffusion |
| TR_orthopi | Orthophosphate-ABC transport |
| TR_pb2 | Lead (Pb^+2^) ABC transporter |
| TR_photon | Photon Transport |
| TR_slf_ABC | Sulfate-ABC transport |
| TR_slf_sym | Sulfate transport in via proton symport |
| TR_urea | Urea transport via facilitate diffusion |
| TR_zn2 | Zinc-ABC transport |
| TR_glycogen | Glycogen Transport |
| TR_cyanophycin | Cyanophycin Transport |

Table S8. **Henry's Constants for Atmospheric Compounds at the Sea-Air Interface** (22)

| **Compound** | **Henry’s Constant (L atm/mol)** |
| --- | --- |
| CO_2_ | 29.41 |
| O_2_ | 769.23 |
| N_2_ | 1639.34 |
